# Is evolution predictable? Quantitative genetics under complex genotype-phenotype maps

**DOI:** 10.1101/578021

**Authors:** Lisandro Milocco, Isaac Salazar-Ciudad

**Affiliations:** Institute of Biotechnology, University of Helsinki, 00014 Helsinki, Finland; Centre de Recerca Matemàtica, 08193 Barcelona, Spain; Genomics, Bioinformatics and Evolution. Departament de Genètica i Microbiologia, Universitat Autònoma de Barcelona, 08193 Barcelona, Spain

## Abstract

A fundamental aim of post-genomic 21st century biology is to understand the genotype-phenotype map (GPM) or how specific genetic variation relates to specific phenotypic variation (1). Quantitative genetics approximates such maps using linear models, and has developed methods to predict the response to selection in a population (2, 3). The other major field of research concerned with the GPM, developmental evolutionary biology or evo-devo (1, 4–6), has found the GPM to be highly nonlinear and complex (4, 7). Here we quantify how the predictions of quantitative genetics are affected by the complex, nonlinear maps found in developmental biology. We combine a realistic development-based GPM model and a population genetics model of recombination, mutation and natural selection. Each individual in the population consists of a genotype and a multi-trait phenotype that arises through the development model. We simulate evolution by applying natural selection on multiple traits per individual. In addition, we estimate the quantitative genetics parameters required to predict the response to selection. We found that the disagreements between predicted and observed responses to selection are common, roughly in a third of generations, and are highly dependent on the traits being selected. These disagreements are systematic and related to the nonlinear nature of the genotype-phenotype map. Our results are a step towards integrating the fields studying the GPM.

## Introduction

A fundamental aim of post-genomic 21st century biology is to understand how genomic variation relates to specific phenotypic variation. This relationship, called the genotype-phenotype map or GPM, is considered by many researchers to be critical factor for understanding phenotypic evolution (1). The GPM determines which phenotypic variation arises from which random genetic variation. Natural selection then acts on that realized phenotypic variation. Thus, natural selection and the GPM jointly determine how traits change over evolutionary time.

Quantitative genetics uses a statistical approach to describe the GPM and predict phenotypic change in evolution by natural selection. This approach has long made significant contributions to plant and animal breeding (2,3). According to the breeder’s equation of quantitative genetics (8, 9), the response to selection in a set of traits 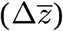, is the product of the matrix of additive genetic variances and covariances between traits (the *G*-matrix) and the selection gradient (*β*), i.e. the direct strength of selection acting on each trait.

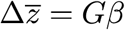

This is the canonical equation for inferring past selection and predicting future responses to selection (10, 11). The *G*-matrix has been interpreted as a measure of genetic constraints on future evolution (12), the diagonal elements of *G* measure the short-term readiness of a trait to respond to selection while the off-diagonal elements measure how a trait evolution is slowed down (or accelerated) by the evolution in other traits.

Developmental evolutionary biology or evo-devo (1, 4-6) is the other main field concerned with the GPM. Evo-devo views the GPM as highly nonlinear and complex (4, 7). Most evo-devo studies, however, do not consider the population level. At that level, it has been suggested that the nonlinearities of the GPM will average out, and hence would not affect the accuracy of quantitative genetics (13), at least in the short-term (14).

Our aim is to study how the predictions of the multivariate breeder’s equation are affected when considering the complex and nonlinear GPMs found in the study of development. For that purpose, we combine a computational GPM model that is based on current understanding of developmental biology (15, 16) and a population genetics model with mutation, recombination, drift and natural selection. The developmental model has been shown to be able to reproduce multivariate morphological variation at the population level (15). It includes a network of gene regulatory interactions and cell and tissue biomechanical interactions. The model’s parameters specify how strong or weak those interactions are. The value of each individual’s parameter are determined additively by many loci. The developmental model produces then, for each individual, a 3D morphological phenotype (see Fig. S1) on which a number of traits are measured: the position of specific morphological landmarks. Individual fitness is calculated as the distance between each of five traits in each individual and these same traits in each simulation’s optimal morphology (see Supplementary Figs. 1, 2). The processes of mutation, development and selection are iterated over generations to simulate trait evolution. At the same time, we estimate *G, P* and *s* in each generation and use the multivariate breeder’s equation to estimate the expected response to selection per generation. We then quantify the difference between expected and observed trait changes in the simulations with a complex development-based GPM (see Fig. S3). The difference we estimate should be regarded as the minimal theoretically possible since we include no environmental noise (i.e. no environmental variance) and *G, P* and *s* are estimated in each generation.

**Fig. 1:**
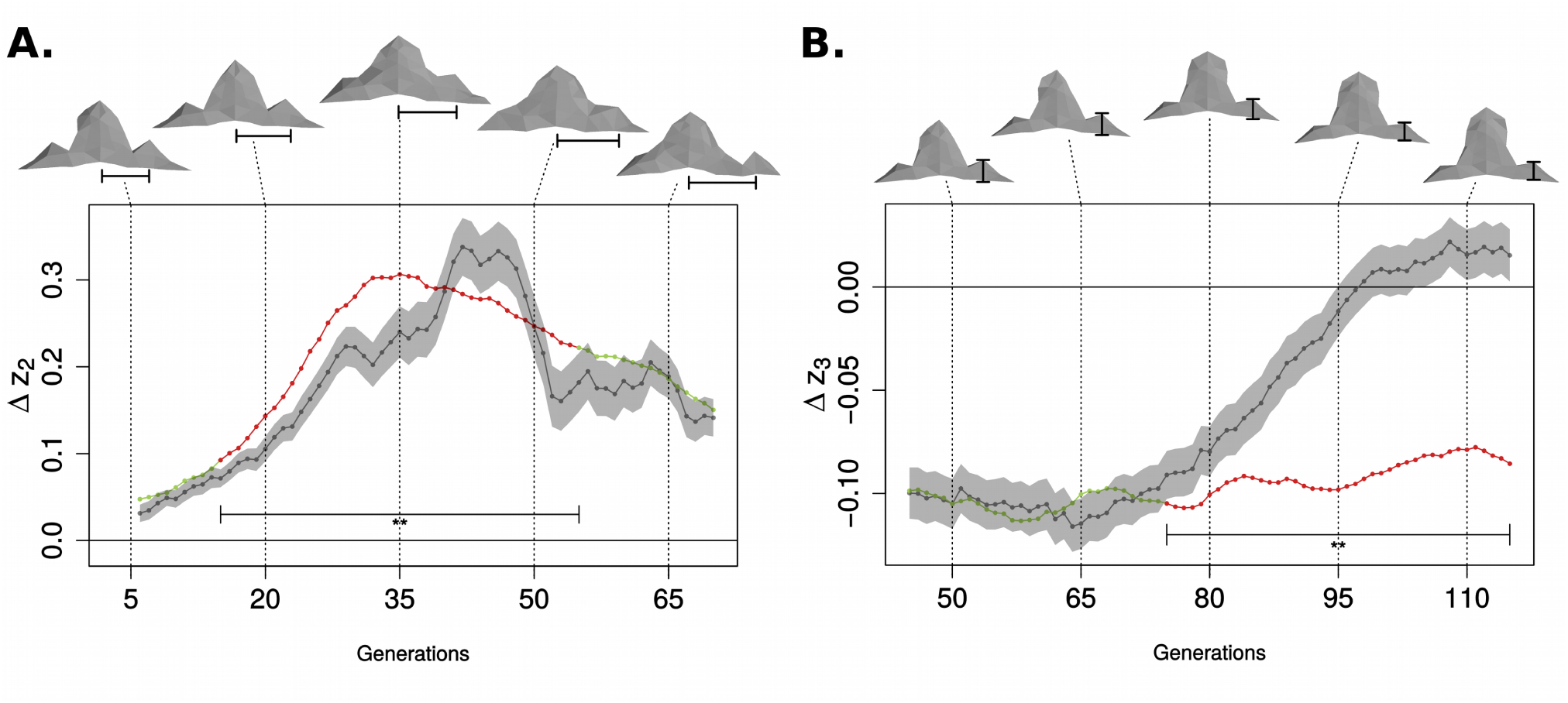
Observed and predicted change can differ in a systematic way over generations. The plots show observed (gray line) and predicted change (green and red lines) for trait 2 in one simulation (**A**) and trait 3 in another simulation (**B**). Predicted change was calculated using the multivariate breeder’s equation and estimates of *G* and *β* in each generation. Observed change is shown with a 99% confidence interval generated from repeating each generation 40 times to minimize the stochastic effect of drift, mutation and recombination. Both changes plotted above are averaged in a time window of 10 generations for more robust estimates (2) (See Materials and Methods). The predicted and observed change are considered to differ significantly when the former is outside of the confidence interval of the latter (marked with a line with two asterisks). Teeth representative of the population at five time points are included above the plots, with a black segment indicating the traits plotted. Evolution for the remaining traits for both simulations is shown in Fig. S4. Panel B shows an extreme case in which a cusp is lost during evolution.

**Fig. 2.**
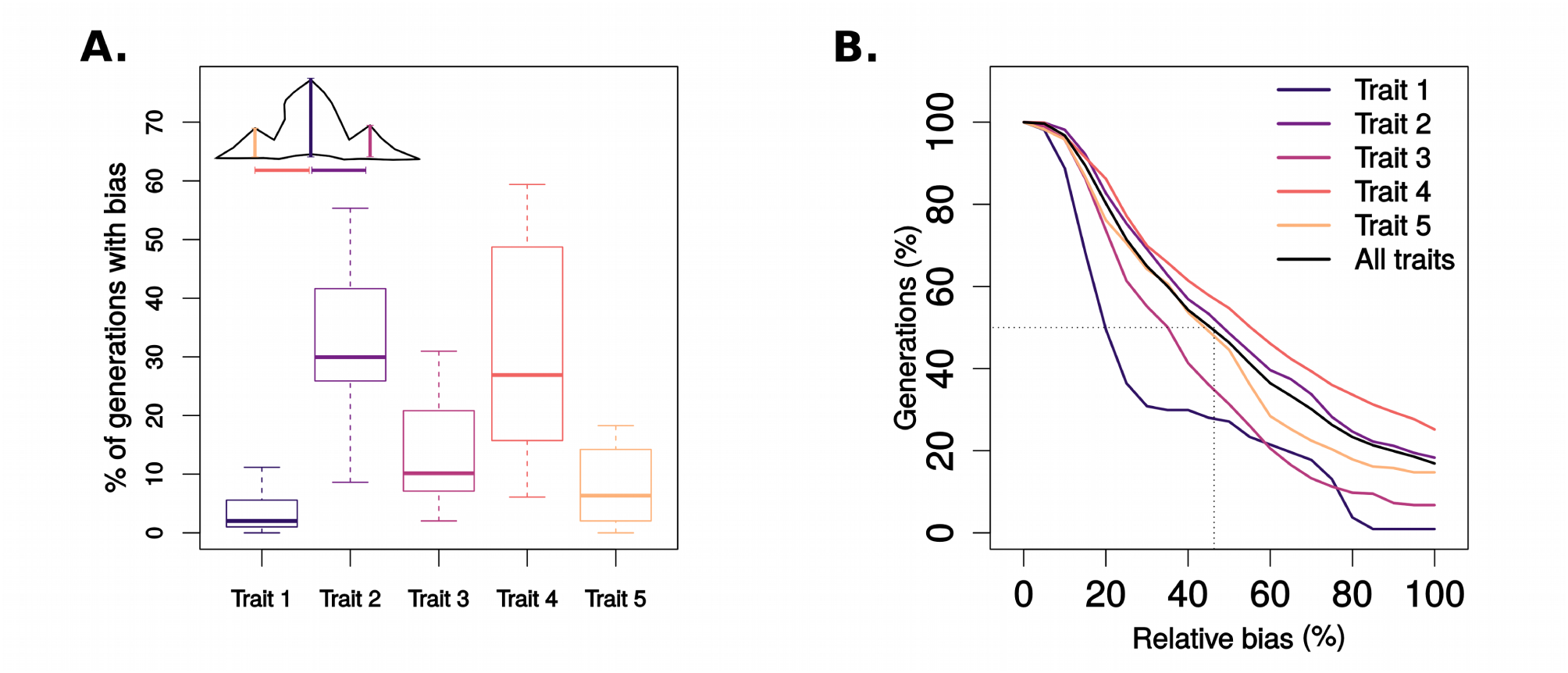
Bias is common and it can be large. Bias is the disagreement between the observed and predicted change in traits mean in a generation. (A) Percentage of generations with significant bias at 99% confidence for all simulations, and for each trait (significance calculated as in Fig. 1, see Materials and Methods). Prediction for change in the mean value of trait 1 rarely shows bias (median of 3.98%) while predictions for trait 2 and 4 are commonly biased (median of 30%). Simulations where the populations goes through regions of the GPM where small parametric changes lead to loss of cusps, such as the one shown in Fig. 1B, where not included. Therefore the plots include data from 14 complete simulations and 197 generations each (13790 data points including all 5 traits.) (B) Percentage of generations with at least a given amount of relative bias (observed change minus predicted change, divided by the observed change). Only generations with a prediction bias with a 99% confidence were used for this plot (a total of 2409 data points out of the total 13790). When considering all traits, half of the generations with significant bias have a relative bias larger than 46%.

## Results

We found that in many generations there is a significant discrepancy, or prediction error, between the trait changes observed in the simulations and the trait changes predicted from the multivariate breeder’s equations (see Fig. 1, Supplementary Video 1, 2, for two example simulations). To study whether this prediction error is merely due to stochastic processes such as drift, mutation or recombination, we re-simulated 40 times each generation of every simulation. Since mutation, recombination and genetic drift are simulated as stochastic processes, different trait changes are observed in each of these 40 simulations (see the 99% confidence intervals shown in a gray shade in Fig. 1). If the multivariate breeder’s equation is accurate, the expected trait changes should not be statistically different from the the mean of these 40 observed changes. However, as can be seen in the simulations in Fig. 1, the deviation is significant for a large number of generations. There is, thus, a systematic prediction error that is not explainable from stochastic processes. We refer to this systematic error as bias. For the simulation shown in Fig. 1A, the bias is large and significant between generations 15 and 55.

Bias was not exclusive to a single trait or simulation (see Fig. S4). Bias was common. Fig. 2 shows the percentage of generations across all simulations that showed significant bias with 99% confidence. For individual simulations the median was that 30% of the generations showed significant bias in traits 2 and 4 (the x-position of each lateral cusps) and less so for the three traits. Bias can be very large, see Fig. 2B. Considering all traits together, the prediction error divided by the observed change per generation is at least 46% for half of the generations exhibiting significant bias.

We found that in our simulations bias arises from nonlinearities in the GPM. Fig. 3 and Fig. S5 show explanatory examples of how specific aspects of the developmental dynamics lead to nonlinearities and then to bias. Bias correlates with measures of the linearity of the GPM around the region of the parameter space where the population is distributed at a given generation (see Fig. S6). These nonlinear regions are quite common and, thus, most populations would pass through one such regions as they evolve.

**Fig. 3:**
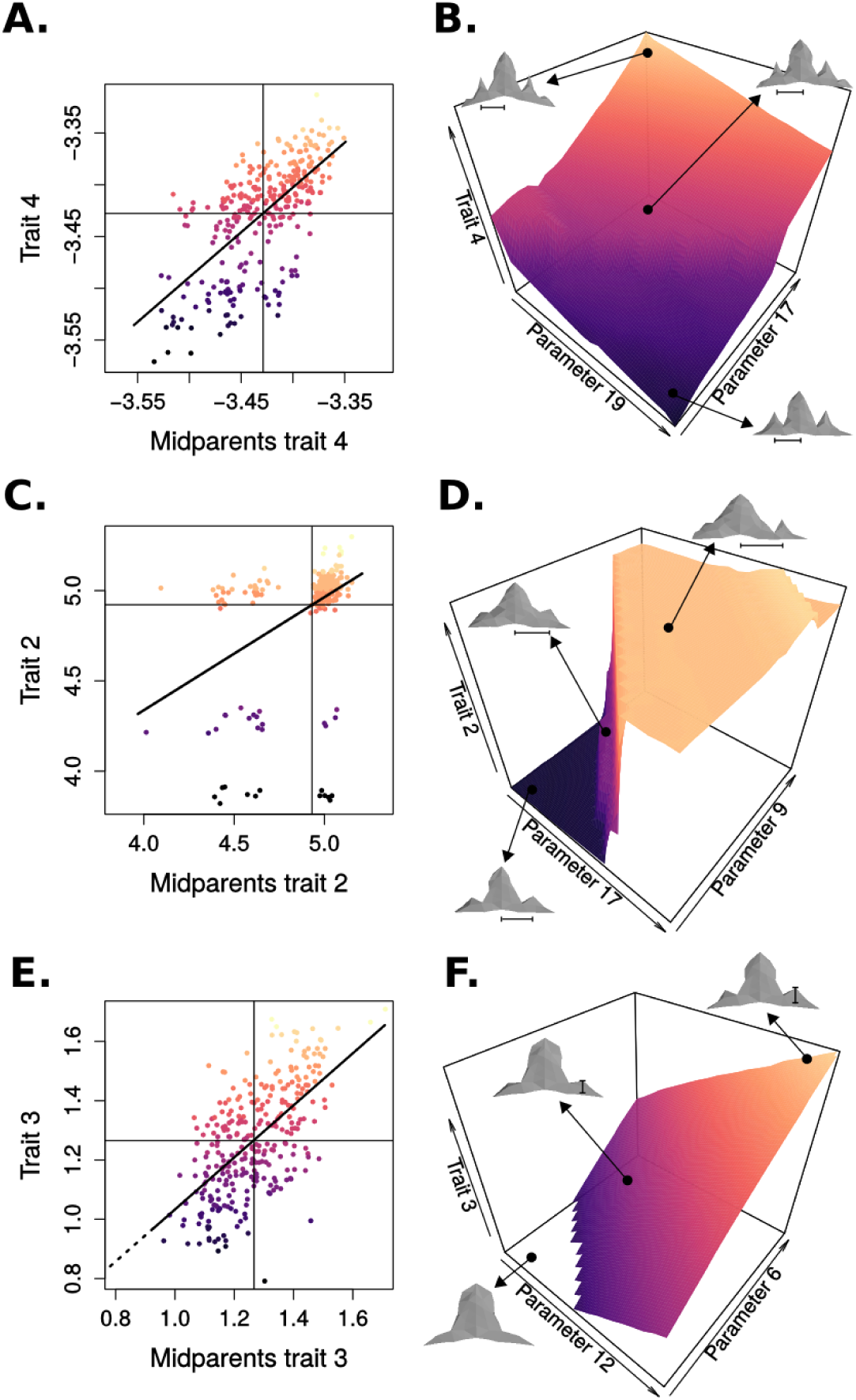
Prediction bias arises when the population is in a nonlinear region of the parameter-phenotype map. The figure shows, for three different generations, the midparent-offspring regression (A, C and E) and a marginal parameter-phenotype (B, D and F) depicting how a trait changes (z-axis) when two specific developmental parameters are varied while keeping all others constant at their population mean value. A and B are for trait 4 in generation 90 of the simulation 110 in Fig. S4. C and D are for trait 2 in generation 50 of the simulation in Fig. 1A. Bias can be explained by the marginal parameter-phenotype maps (B-D), showing that small changes in parameter values in this region produces relatively large changes in the measured traits. E and F correspond to trait 3 in generation 100 of simulation 107 in Fig. 1B. The marginal parameter-phenotype map (F) shows that, when both parameters are low, the phenotypes produced lack lateral cusps. Bias arises because the linear approach of multivariate breeder’s equation estimates that teeth with smaller values for trait 3 can be produced when in fact they cannot. For panels A, C and E 300 random individuals are plotted as points.

The bias we encountered is not specific of our development model. We have ran also a model not based on development, in which traits were simple mathematical functions of some arbitrary model parameters (see Supplementary Information). When a linear map was used, no prediction bias was found. In contrast, nonlinear GPMs lead to bias as with our development model (Fig. S7).

Population parameters such as population size, mutation rate, selection strength and number of loci had a modest effect on bias (Fig. 4). They affect how fast the population moves in the trait and developmental parameter space and then, indirectly, the likelihood of encountering a nonlinear region of the parameter space leading to bias (see Fig. 4 and Supplementary Information).

**Fig. 4:**
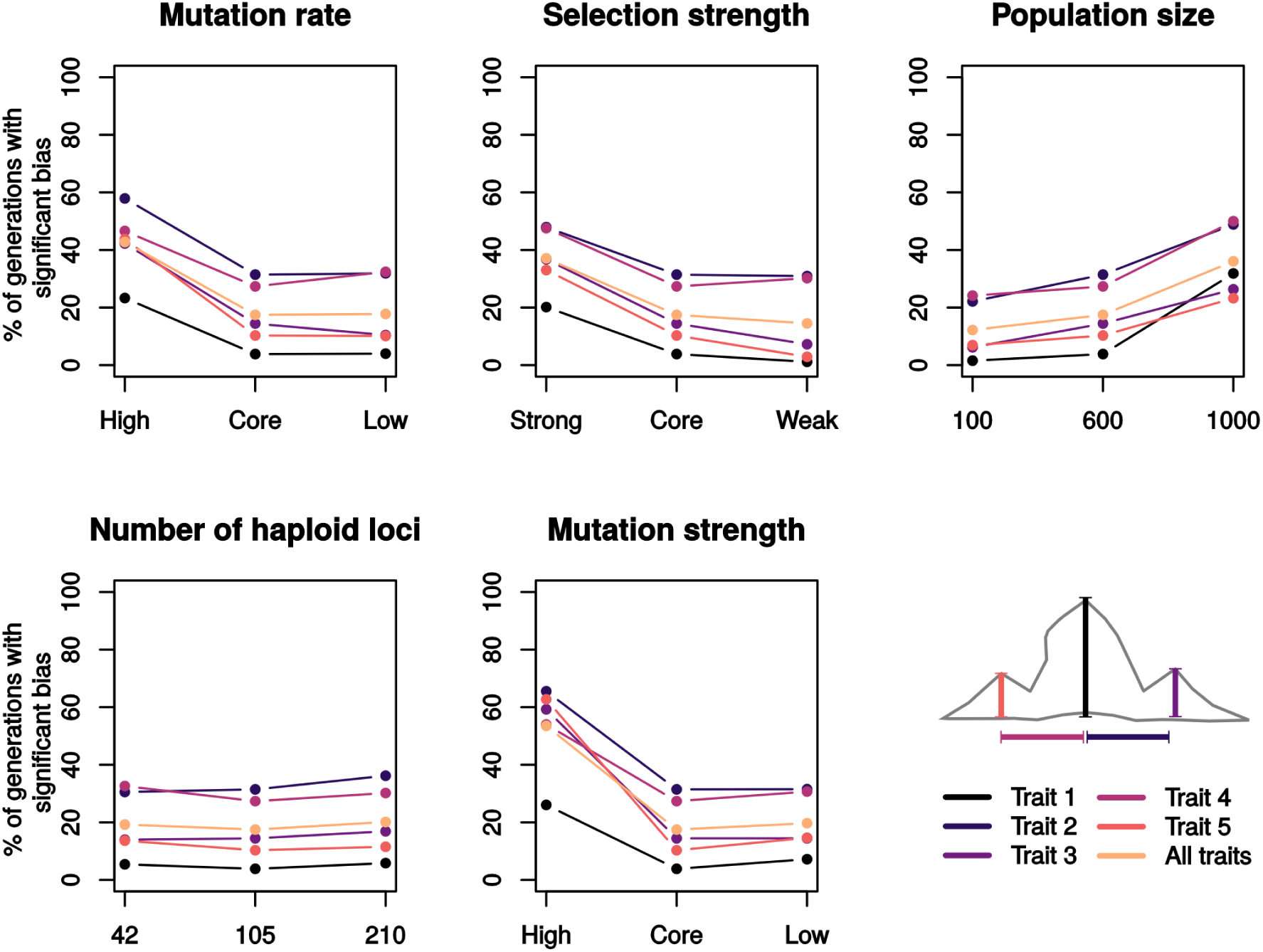
Population genetic parameter have only a modest effect on the bias. The plots above show the median percentages of generations showing significant prediction bias (99% confidence) for each trait, when modifying one population genetics parameter from the core parameter set, keeping the others to the core value (14 simulations per point in the plot, excluding the ones where there is loss of cusp as shown in Fig. 1B). Parameter values are shown in Table S1. In general, our exploration shows a saturating relationship between the evolutionary parameters and the prediction bias. This is, there is a modest dependency of the bias with the parameters but the amount of bias reaches a plateau, determined by the characteristic of the GPM which are not dependent of population-level parameters. A detailed explanation of the effect of each parameter on the bias is given in the Supplementary Information (*The effect of population parameters on the bias.)*

## Discussion

Bias can be understood better by considering how additive genetic variances are estimated. Although there are several ways to do so (see Materials and Methods), they all assume a linear relationship between traits and the genetic relatedness between individuals. This assumption does not always hold in our development-based model, which means that the linear transformation used in breeder’s equation can significantly over-or underestimate the change in the mean of traits. The examples in Fig. 3 and Fig. S8 show, for three generations with bias, the parent-offspring regression for a trait, and the region of the developmental parameter occupied by the population. In this region, small genetic changes lead to small changes in the developmental parameters but these lead to relatively large changes in some traits. As a result, when the population crosses this region of the developmental parameter space on its way to the optimum, the change in some of the traits is different to that predicted from the linear approximation of the multivariate breeder’s equation.

Even when variation no variation is observed beyond some critical trait value bias can still occur (Figure 3E). Some combinations of traits values, i.e. some morphologies, can not arise in the development model, at least not without dramatic changes in the developmental parameters. As a result, it can sometimes occur that a trait can not evolve beyond the critical trait value even if there is a selective pressure for it. In this situation bias arises because, even if there is no variation above or below the critical trait value, there is trait variation before the population reaches this value. Because this variation, the parent-offspring regressions for that trait (and between this trait and the others) are non-zero; see Figure S9. Based on these regressions the multivariate breeder’s equation estimates that variation beyond this critical trait value should exist, and thus a possible response to selection, when in fact it does not (see Figure S9). This prediction bias can persist over time because even if the population is pushed by selection close to the critical trait value, mutation and recombination spread it over a range of values before the threshold and, thus, a non-zero parent-offspring regression persists (see Figure S9B).

There is already some theoretical literature stating that the assumptions of the breeders equations may not always hold. Most previous studies, however, do not directly consider the GPM (17, 18). An exception is Rice’s formalization of adaptation under known and nonlinear GPMs (19). This previous work, however, does not explain how nonlinear GPMs may actually be. Our study quantifies to which extent the predictions of breeder’s equations hold for realistic development-based GPMs as we currently understand them. Compared to theoretical evolutionary genetics models (20, 21), our model does not assume a specific pattern of epistasis among loci, instead, such patterns arise from the dynamics of the development model and are not compatible with these assumed patterns (see Fig. S10 and Supplementary Information).

Although our development model is based on tooth morphogenesis, there is no reason to expect that the complexity of the GPM should be very different for other organs of a similar morphological complexity (22, 23). In fact, in the supplementary information we explain how even arbitrary GPMs, as long as they are nonlinear, would produce similar biases (see Fig. S7). In addition, most, if not all, mathematical models of development or gene network dynamics are highly nonlinear and lead to complex GPMs similar to those in our model (23, 24).

The prevailing view in quantitative genetics is that breeder’s equations is very accurate when applied to single traits (3). For selection for multiple traits at the same time or when selecting for some traits and studying the correlated response in other traits, however, the scarce empirical evidence does not always fit the expectations from breeder’s equation (3, 25-27). Our results suggest a simple reason for these inaccuracies and provide a theoretical nonlinear-GPM perspective on what to expect when multiple trait selection experiments finally become more common.

There is a long ongoing apparent discrepancy between evolutionary developmental biology and quantitative genetics (7, 9, 12, 14). One approach views the GPM as simple enough, at least in practice, for trait evolution to be predictable from linear statistical approaches. The other views the GPM as highly nonlinear. Our results present a potential point of connection between these two views. The predictions of quantitative genetics would often work accurately but, very often too, they would show relatively large prediction errors that can not be corrected by better estimating quantitative genetic parameters since they are due to the way the nonlinearity of the GPM.

## Materials and Methods

In this work, evolution is modeled using an individual-based algorithm similar to that in (16). The complete model is composed of a developmental model and a population model. The developmental model is used to generate each individual’s phenotype from its genotype. The population model is used to decide the genotypes in each generation based on genotypes from the previous generation through selection, mutation, drift and recombination. Each model has a set of parameters, the developmental and populational parameters respectively. Both the population and developmental model were written in Fortran 90. The data from the simulations was analyzed and visualized using R. In addition, in each generation, the matrix of additive genetic variances and covariances between traits, *G*, the matrix of phenotypic variances and covariances between traits, *P*, and the selection differentials, *s*, are estimated. From that an expected response to selection is calculated from the multivariate breeder’s equation and compared with the one observed in the simulations for the next generation.

### Developmental model

The developmental model used for our simulations is a computational model of tooth development (15). This model provides an example of a genotype–phenotype map for the morphology of a complex organ. The tooth developmental model is a mathematical representation of the current understanding of the basic gene network behind tooth development, and includes the basic cell behaviors (cell division and cell adhesion), cell mechanical interactions and their regulation by gene products known to be involved in the process. The model is mechanistic in the sense that from this hypothesis and some very simple initial conditions-i.e. a flat epithelium representing the initiation of tooth development-the model reproduces how the morphology and patterns of gene expression in 3D change during development until specific adult tooth morphologies are reached. The dynamics of the model are determined by the value of a set of developmental parameters that specify the magnitudes of several biological and physical properties of the cells, tissues and molecules involved in the process such as, proliferation rate, molecular diffusion rates and regulation interactions between genes. The model’s output is the three-dimensional position of tooth cells and, thus, tooth morphology. For a more detailed description of the model we refer the reader to the original publication introducing it (15). The tooth developmental model is suitable for our question because it is able to produce realistic population-level phenotypic variation (15). Furthermore, it is based on developmental biology and therefore includes epigenetic biophysical factors, such as extracellular cell signaling and diffusion in space and mechanical interactions between cells, which have been extensively proposed to lead to complex genotype-phenotype maps (28, 29).

For our study, measurable morphological traits needed to be defined in the tooth shape to be tracked through the generations of the population. The phenotypic traits under study were defined to be the spatial position of landmarks in the tooth morphology. Landmarks were located in the three tallest cusps arising in each tooth. The three tallest cusps are the first three cusps to be formed during development (30). Fig. S1 shows the location of the three landmarks in an example morphology. Traits 2 and 4 are the x-positions of the posterior and anterior cusp, respectively. Traits 1, 3 and 5 are the heights of the central, posterior and anterior cusps. Height is measured from the cusp to the baseline of the tooth. The source code and executable files for the model can be found in https://github.com/millisan/teeth-evo

### Population model

The developmental model was embedded in a population model. Table S1 shows the populational parameters. We defined a core set of population parameters and studied deviations from those core values, other population parameter sets. For each each parameter set, 20 simulations each of 200 generations were ran using different optima to explore the behavior of the system.

The population model considers 4 steps per generation: 1) a mapping between genotypes and developmental parameters; 2) a mapping between these parameters and phenotypes (the developmental model); 3) a mapping between phenotypes and fitness; 4) reproduction (with recombination) and mutation on the genotypes. By iterating steps 1 to 4 in each generation, we simulated how the genotypes and phenotypes of the population change over generations (Fig. S3).

#### 1) Mapping between genotypes and developmental parameters

The input of the developmental model are the values of the developmental parameters. Each of these parameters corresponds to a developmentally relevant interaction, but not necessarily to a single gene (see (15)).

Each individual’s genotype is diploid with a fixed number of loci. Each allele in a loci has a specific quantitative value, its allelic effect. The developmental parameters values are determined as the sum of the contributions of a fixed number of loci (similar to (31)). We purposely use this very simple yet unrealistic mapping because our focus is on how the part of the developmental dynamics we understand the better (i.e the developmental model) affects phenotypic variation and evolution, without confounding effects. There is no pleiotropy since in the mapping from genes to parameters each loci contributes to the value of one and only one developmental parameter. Pleiotropy from genes to phenotype arises from the developmental dynamics which map parameters to phenotypes.

#### 2) Mapping between developmental parameters and phenotypes

This is accomplished by running the developmental model, for each combination of parameters corresponding to each individual To determine the trait values, each tooth is first centered so that the main cusp is located at x=0. The z component of each landmark is determined as the distance on the z axis from the landmark to the baseline of the tooth (see Fig. S1; as in (15))

#### 3) Mapping between phenotypes and fitness

Each simulation had a different optimal morphology. For each population parameter set, a total of 20 optima were chosen (see Fig. S2). Each of the 5 morphological traits was chose to increase or decrease in respect to an initial reference morphology (see the “Initial population” section). This leads to a total of 2^5^ = 32 trait combinations. This number reduces to 20 when removing one of two mirror symmetric combinations. Because we wanted selection to act with the same strength on every trait, the optimum value for each trait was set at an equal distance from its mean value in the starting population. The optimum morphology was then determined by adding or subtracting 3 units to the mean value for the trait in the population at generation 1, depending if selection was upwards or downwards, respectively.

In each generation, the fitness of each individual in the population was calculated as a function of the distance of its trait values and the optimal trait values as

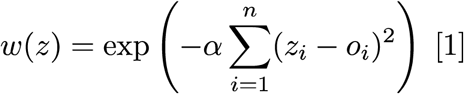

Where *z* is the vector of *n* trait values for the individual, *z*_*i*_ is the *i*^*th*^ element of that vector, *o*_*i*_ is the *i*^*th*^ element of the vector of optimum trait values and *w* is the fitness function. The steepness of the selection gradient is determined by the parameter *α*(see Table S1).

The parents of each individual in a generation were chosen at random from the individuals of the previous generation. For each individual in the previous generation, the probability of being chosen as a parent was equal to its fitness divided by the sum of the fitness of all individuals in the population in that generation. In this way we implement both natural selection and genetic drift.

#### 4) Reproduction, recombination and mutation

Once the parents are selected, they produce one gamete each by randomly selecting one of the two alleles for each loci with equal probability. The gametes of the parents then fuse to form a diploid genome. In the gamete formation process, there is a per-loci probability of mutation, parameter *muta*. Mutation is implemented by adding a random number to the mutating loci, drawn from a normal distribution with a zero mean and *τ* standard deviation. We call *τ* the mutation strength.

#### 5) Initial population

The population has fixed size. The initial population is the same in all simulations with the same parameter set. Each locus starts with ten equally frequent alleles with allelic effects drawn from a normal distribution. The mean of such distribution was such that when summing over all loci one obtains the reference developmental parameter values, i.e. the developmental parameter values that reproduce the morphology of the postcanine tooth of the ringed seal (*Phoca hispida ladogensis*). The variance of the distribution of genetic values was set equal to *τ* (see *4 Reproduction, recombination and mutation*). These genetic values were then used in simulations under stabilizing selection for 100 generations. Stabilizing selection was performed by using the mean traits of the population at the first generation as optimal morphology (see *3 Mapping between phenotypes and fitness*). The resulting population was used in the actual simulation experiments as the initial condition.

### Quantitative genetic model

What we call the observed change is the change in trait means in the simulated population between two consecutive generations. This vector is to be compared with the predicted change using the multivariate breeder’s equation on the *G, P* and *s* estimated from each generation in the simulation,

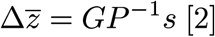

Where 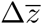 is the between-generation change of trait means, *G* is the matrix of additive genetic variances and covariances, *P* is the matrix of phenotypic variances and covariances and *s* is the selection differential. It is known that the parameters of the breeder’s equation can change in time, although there is debate regarding how fast this occurs (33-37). To avoid errors arising from outdated estimations, *G, P* and *s* were calculated for each generation. Note that *P*^-1^ *s* = *β*, where *β* is the selection gradient i.e. the direct strength of selection acting on each trait.

*G* was estimated in two ways. The first was directly by midparent-offspring regression (2). Animal models were also fitted to the data. In each generation, a nested design was generated from the population. 100 individuals were randomly selected as sires and 200 as dams. Each sire was mated with 2 dams, and 2 offspring were generated from each couple. The nested design allowed to have phenotypic information on full-sibs, half-sibs and parents. The data was fitted with an animal model and restricted maximum likelihood (REML) estimates of additive genetic variance were obtained using the software WOMBAT (38). The animal model used was the simplest possible,

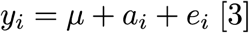

With *y*_*i*_ the vector of traits for the *i*^*th*^ individual, *a*_*i*_ the breeding value and *e*_*i*_ the residual term. The variance-covariance matrix for additive genetic values estimated from midparent-offspring regression and the variance-covariance matrix of the traits were used as starting estimates for the random effects. Both methods yielded similar estimates of *G*.

The selection differential *s* was calculated, in each generation, as the covariance between relative fitness and each trait value (8).

### Statistical Analysis

Due to the stochastic nature of the evolutionary process, we had to develop ways to reveal the deterministic part of the difference between observed trait change and the trait change predicted from the multivariate breeder’s equations. To do this, we performed repetition simulations. These consist in re-simulating the same generational step multiple times. In each repetition, stochastic processes such as recombination and mutation differ, leading to a stochastic distribution of observed changes, and therefore a distribution of errors. This approach allows to statistically test against the null hypothesis “the mean predicted change equals the mean observed change”. Rejecting the null hypothesis implies that there is a systematic difference between the measurement and the prediction that cannot be attributed to randomness. We refer to such systematic component of the error as bias.

For each repetition we can calculate an observed change as the difference in means between the original population and the one generated in the repetition. We then compare this distribution of observed changes with the predicted change using the breeder’s equation. For all parameter combinations studied, and for all 200 generations of the 20 simulation in each of these combinations, we performed 40 repetitions. This leads to a total of 1760000 simulated generations.

Because we are working with a finite population the effect of sampling is not totally eliminated even with this approach. This is why we average the observed and predicted change in a time window of 10 generations. This is equivalent to doing a regression of the response to selection against generations, as typically done in artificial selection experiments (2). We claim there is significant bias when the predicted change deviates from the distribution of observed changes by more than 3 standard errors of the observed mean (this is the 99% confidence in a normal distribution, see Fig. S1).

## Acknowledgments

We thank Tobias Uller, Asko Mäki-Tanila, Pascal Hagolani, Jukka Jernvall, Juha Merilä, Heikki Helänterä, Mihaela Pavcilev, Roland Zimm, Renske Vroomans and Jhon Alves for useful comments.

## Funding

This research was funded by the Finnish Academy to ISC (315740 and 272280) and ILS to LM.

## Authors contributions

Funding acquisition and Supervision: ISC, Conceptualization and Writing: ISC and LM. Investigation, Software, Methodology: mostly LM but also ISC, Formal analysis, Validation and Visualization: LM.

## Competing interests

The authors declare no competing interests.

## Data and materials availability

All data is available in the manuscript or the supplementary information. Data and computer code used available at https://github.com/millisan/teeth-evo

## I Supplementary results

To further confirm that prediction bias arises from the nonlinearities of the parameter-genotype map, we performed evolutionary simulations where the GPM was defined by simple mathematical explicit functions. We compare a linear and a nonlinear function. We tested the predictions of the multivariate breeder’s equation for both functions while keeping all other evolutionary parameters constant and equal to the core parameter set (see Materials and Methods).

For both mappings, 4 developmental parameters were used and 2 traits were defined. Traits are symbolized as Φ and developmental parameters as *u* (following the notation from Rice, 2002).

The linear map used was:

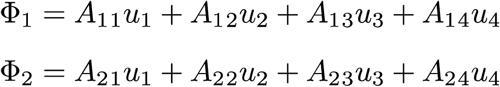

The nonlinear map used was:

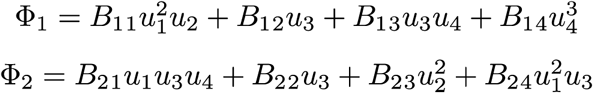

Matrices *A* and *B* were randomly generated from a normal distribution with mean 0 and variance 1.The exact values of these matrices had no effect on our results. The evolutionary algorithm is exactly the same as described in the Materials and Methods section, with only the GPM being different. Briefly, each individual’s genotype is diploid with a fixed number of loci, and is stored as a list of genetic values (see Materials and Methods). The developmental parameter values are calculated as the sum of a fixed number of loci that contribute to the parameter. Each locus starts with ten equally frequent alleles with allelic effects drawn from a normal distribution with mean of 0 and variance equal to the per-loci mutational variance. These genetic values were then used in simulations under stabilizing selection for 100 generations. The genotypes after those generations were used as initial conditions to the actual simulation runs.

A total of 30 simulations with random optima were ran for each map, linear and nonlinear, for 200 generations. Every generations, prediction bias was estimated by re-running the same generational step 40 times, allowing to test against the null hypothesis “the mean observed change equals the predicted change”.

The percentage of generations that showed bias with 99% significance are shown in Fig. S7A. All optima and generations were used for the plot. For the linear traits, a median of 1.57% of generations showed bias. This is negligible and can be attributed to estimations error of the G matrix due to finite population size, a phenomenon known as the Turelli effect (Turelli, 1988; Jones *et al.*, 2004). Simulations with the nonlinear map on the other hand show a median of 32% of generations with significant bias.

Fig. S7B shows the relationship between local nonlinearity of the parameter-phenotype map and the observed bias, analogous to that shown in Extended Data Fig 6. Only data from simulations using the nonlinear map were included in this plot since simulations with linear map gave nonlinearities of the order of 10^-16^, with median error of 0.002155379. As in Fig. S6, it can be seen that both the spread and magnitude of the bias increase with the nonlinearity of the parameter-phenotype map. Note that this measure of nonlineairty includes both monotonous and nonmonotonous parameter-phenotype maps, sensu Gjuvsland and collaborators (Gjuvsland *et al*, 2013a).

## II Supplementary discussion

In this supplement we discuss how our work relates to previous studies and how our approach and findings are different. Previous research includes general criticisms to the multivariate breeder’s equation approach, models considering the inadequacy of the infinitesimal model, the non-normality of trait distributions, epistasis and nonlinear GPMs, or a combination of these issues.

One great advantage of the multivariate breeder’s equation is that it can be used for any phenotype without requiring any knowledge of its genetics or development. Only selection, the trait values in individuals and the genetic relatedness between individuals in a population need to be known. This simplicity, however, comes at the cost of assumptions that may often not hold up.

### General criticisms

The multivariate breeder’s equation has been found to be inaccurate in some of the few phenotypes in which observed and predicted changes have been compared (see Roff 2007 for a review). Inaccurate measurement of either selection or genetic variance, changing environmental conditions, or an incomplete picture of all traits under selection can lead to large inaccuracies in the predictions provided by the multivariate breeder’s equation (Merilä *et al*. 2001; Barton & Turelli 1989, Hadfield 2008, Wilson *et al*. 2006; Morrisey *et al.* 2010).

From another perspective, Pigliucci (2006) points out technical problems such as the arbitrariness in defining the traits included in the G matrix and the artificiality in the breeding programs commonly used to estimate G, usually carried out with lab-maintained populations outside their natural conditions. The author also criticizes the G-matrix research program on conceptual grounds, such as the exclusively local value of G (only valid for a given population at a given time point), the highly degenerate mapping between G-matrices and underlying genetic and developmental structures, and the difficulty of discriminating the effects of selection and drift from G structure.

Much work in developmental evolutionary biology is implicitly or explicitly critical with quantitative genetics in general and with the breeder’s equations in particular (Alberch, 1982; Müller, 2007; Müller, 2017). These critiques are rooted in the fact that the study of development has consistently shown that the GPM is rather complex and nonlinear (Alberch, 1982; Polly, 2008). From this perspective the linear approach of breeder’s equations is seen as a gross oversimplification, especially at the macro-evolutionary scale.

Polly (2008) points out three key consequences that developmental interactions can have on morphological evolution and that evolutionary quantitative genetics cannot fully account for. We found evidence of all three points in our simulations. Firstly, even with continuous variation at the level of underlying developmental parameters, traits may occasionally exhibit small jumps in evolution (see the jump in the position of the posterior cusp in Fig. 1A.). It is important to notice, as we explain in the main text discussion, that these jumps are rather small, of at most 10% of the trait value, and, thus, may not be perceived as trait discontinuities, especially given the environmental effects and measurement errors always present in experiments. Secondly, Polly argues that developmental interactions may limit production of certain phenotypic variants in non-additive ways, leading to a bias in the prediction of the response to selection. This is mirrored in our findings of the prediction bias in certain regions of the GPM. We explicitly show that the structure of additive genetic variances and covariances in the *G*-matrix does not always summarize all associations between traits, leading to the bias in the prediction. Lastly, Polly mentions that some nonlinearities will lead to loss or appearance of structures. Evidence for this can also be found in our simulations and is associated with problems in the application of the breeder’s equations (see Fig. 1B and associated problems when the population is in a region of the GPM where a small change in parameter values leads to loss of the posterior cusp).

### The infinitesimal model

One founding assumption of quantitative genetics is the infinitesimal model (Fisher, 1918). According to this model, trait values are determined by a large number of loci of small and fixed effect. By fixed effect we mean an effect that is independent of that of other loci. This assumption does not seem to hold for the traits for which we know something about the underlying loci. First, there is plenty of evidence for alleles of relatively large effect and, in fact, for a diversity of magnitudes of effects (Alberch, 1982; Gibson, 1999; Eyre-Walker and Keightley, 2007). Second, as explained through this article and elsewhere, everything we know about development indicates that genes can construct the phenotype and, thus, contribute to their variation because they interact with each other (Waddington, 1957, Wright, 1978, Alberch, 1982, Rice, 2002; Salazar-Ciudad 2006; Uller *et al.* 2018). This strongly suggests that the effects of loci are not fixed, but depend on each other, i.e they are epistatic.

In the context of quantitative and population genetics, epistasis between a set of loci is only relevant if these loci are polymorphic. In other words, only variable loci can contribute to trait variation and, thus, have a role in the prediction of the response to selection. When this occurs there is *statistical* epistasis (Cheverud and Routman 1995, Álvarez-Castro and Carlborg 2007, Hansen 2008, 2013). That two gene products interact, for example, during development, is called *functional* epistasis (Álvarez-Castro and Carlborg 2007, Hansen 2013 and *physiological* in Cheverud and Routman 1995). That two genes interact during development does not imply that they are statistically epistatic to each other since they may not be polymorphic. This fact has been used (Hill *et al.*, 2008; Hansen, 2008) to argue that the understanding of how genes interact during development, and the resulting nonlinear GPM, is not necessarily relevant for quantitative genetics. This has been taken to imply that, given the general success of quantitative genetics, the infinitesimal model is a good approximation (Hill *et al.*, 2008; Turelli and Barton 1994). As argued in the main text, however, this success is only evident for the case of single traits. In addition, since the vast majority of genes interact with many other genes and allelic variation is common (Wright, 1978), one should expect epistasis to be relevant for the quantitative genetics of most traits. In other words, functional epistasis should often lead to *statistical* epistasis. This implies that the effects of most loci would not be fixed and, that thus, the infinitesimal model would be hardly tenable. Others, however, argue that either functional epistasis is not so prevalent or that loci are not so polymorphic to make statistical epistasis relevant enough to invalidate the infinitesimal model (Hill *et al.*, 2008). Overall there is a long-lasting controversy about the infinitesimal model and epistasis. The emerging consensus view seems to be that the infinitesimal model is, at best, a convenient idealization that may roughly hold in a number of situations but not in others (Huang and Mackay, 2016). The idealized nature of the infinitesimal model could underlie the so-called missing heritability problem. This is basically the empirical observation that the total sum of the single-locus variation of alleles affecting a given trait accounts for only a small fraction of the total heritability of the trait (Zuk *et al.*, 2012).

Strictly speaking the infinitesimal model is not required for some of the inferences made in quantitative genetics. The infinitesimal model was central to quantitative genetics because, originally, quantitative genetics was seen as an extension of multilocus population genetics. Continuous traits were then assumed to be affected by many loci, each of small, fixed and additive effect. This allowed to define things such as additive genetic variance and heritability based on loci effects. This is not possible, however, if the infinitesimal model does not hold. In this case, one can still do quantitative genetics, simply one should treat quantitative genetics as a first order approximation to phenotypic change based on the parent-offspring regression (Rice 2004, 2012). In this case the breeders equations should be interpreted as the linear prediction one can cast on the response to selection based on the regression between parents and offspring (or based on covariances in the case of the multivariate breeder’s equation).

### Models with epistasis

There have been attempts to develop quantitative genetic frameworks that can accommodate for certain types of epistasis. Hansen and Wagner (2001) introduced a multilinear model in which the effect of an allele substitution in a locus is modelled as a linear combination of locus effects and pairwise epistatic effects among loci. Using this description, the authors develop expressions for usual quantitative genetics quantities such as additive genetic variance, and most relevant to us, a modification of the breeder’s equation that takes into account epistasis. Carter *et al* 2005 completed this analytical work with simulation studies using the multilinear model, and showed that directional epistasis modifies the response to selection from that expected by the breeder’s equation.

A distinctive feature of our work is that we simulate each individual in the population using a mechanistic model of development. The dynamics of this model cannot be framed in terms of the multilinear model. The limiting assumption in the multilinear model is that changing genetic background of a locus causes a scaling of the effects of all substitutions in that locus by the same scale factor. This is not the general case when using the tooth model, as shown in Exteded Data Fig. 9. The equations developed in Hansen and Wagner (2001) for predicting the response to selection are therefore not applicable to our system or to any GPM without this specific scaling.

Our model does not have the limitation of the multilinear model but it is not analytically tractable. Indeed, while working in the multilinear model, Carter *et al* (2005) hints towards the importance of the work we are presenting here to the understanding of the effect of nonlinear interactions on the response to selection: “To study truly nonlinear forms of gene interaction, we have to turn to highly specific architectures, and rely almost exclusively on computer simulations.”

### Non-normal distributions

Much theoretical work has been done to study the effects of the departure from the assumption of normality in the distribution of traits. Among the first to do so were Turelli and Barton (1990, 1994). Most notably, they derived an expression for the deviation in the expected response to selection when breeding values are not normally distributed, under the assumption of additivity among loci (i.e. each loci contributing a fixed amount to the final phenotype, so that the phenotype is the sum of each individual loci’s contribution). More recently, Bonamour 2017 used the equations derived in Turelli and Barton 1990 and computer simulations to study the effect of skewness (asymmetry) in phenotypic distributions on the measurements of selection and on the response to selection.

Turelli and Barton 1994 conclude that the infinitesimal model is robust to deviations from normality. The conclusion is, however, only valid under the assumption of additivity. Departure from normality in phenotypic distribution in their analysis therefore does not arise from nonlinear genetic interactions but other effects such as environment, recombination and linkage disequilibrium. Their approach is, thus, fundamentally different to the one presented in this paper, where the focus is in the GPM itself.

### Models with nonlinearities

Rice’s work (2004) shows that breeder’s equations assumes the local linearity of GPM and that the predictions of the multivariate breeder’s equation could be inaccurate under complex nonlinear GPMs. This work demonstrates that this could happen, but does not include an understanding of actual GPMs from which to evaluate how large and frequent should these inaccuracies be.

Rice (2004) developed a mathematical population framework to study the dynamics of adaptation under arbitrary GPMs, including nonlinear ones. This framework requires no assumptions about the interactions between loci nor about trait distributions. This framework, however, assumes that the GPM is known. Based on the derivatives of the traits with respect to some underlying factors (e.g. genes or developmental parameters) and the moments of the trait distribution, Rice develops formulae to predict the response to selection.

Morrissey and collaborators (Morrissey 2015, de Villemereuil 2016) use mathematical functions to map normally distributed genetic and environmental factors to traits, similarly to Rice. Classical quantitative genetics can be applied at the level of the underlying factors and then the results can be translated back to the traits by using the functions. This allows to estimate classical parameters such as additive genetic variance of the traits. The authors also developed a predictive equation for the response to selection for nonlinear GPMs. As in the case of Rice work this mathematical description of the GPM does not arise from the theory itself but needs to be provided from elsewhere. Our approach is intrinsically different: it has a theory on how the GPM is and then explores its evolutionary consequences. It is important to notice that our theory of the GPM is coming from a theory of development and, thus, it is based on a completely different sort of evidence and logic than Rice’s work, models with epistasis and quantitative genetics in general.

In addition, the function used Morrissey’s work is assumed to be invertible. This restricts the application of the method to one-to-one mappings between genetic and phenotypic variation. This is problematic since most GPMs are known to exhibit many-to-one mappings (Wagner, 2011), including the one of the tooth model (Salazar-Ciudad and Marin-Riera, 2013).

Using Morrissey’s approach, De Villemereuil (2016) shows that systematic biases in the predictions of the breeder’s equation can be encountered when the mapping function used is logarithmic. This prediction bias is similar to the one we describe in this work. In here, however, we do not impose a function mapping parameters to phenotypes. Rather, the mapping arises from the dynamics of the developmental model and changes with time. Because the mapping from inputs to traits changes with time, we observe dynamics in which prediction bias changes both quantitatively and qualitatively, even finding regions where there is no prediction error, despite the developmental function being nonlinear (see Exteded Data Figure 6, where high nonlinearities can be associated with negligible bias.)

The approaches just described do not aim to understand the GPM, they aim to understand how the GPM may affect trait adaptation. In these approaches the GPM is given as an explicitly defined function between genotype, or other underlying factors, and traits. Many mathematical approaches that aim to understand the GPM use a quite different approach that models the network of basic gene product and cell interactions involved in the production of the phenotype, such as in development or physiology. As a result of this modelling of the “microscopic” rules a specific relationship between genetic and trait variation arises, a GPM.

Gjuvsland and collaborators (Gjuvsland 2011, Gjuvsland 2013a, Gjuvsland 2013b) take one such alternative approach. They study the adequacy and predictive value of some central concepts in quantitative genetics in the context of a realistic nonlinear GPM. In that sense their approach is similar to ours, they simply address a different set of questions and use simpler models based on physiology, rather than on development, and deal with single traits. As in the current work, they emphasize the importance of describing the GPM as arising from dynamic physiological systems. They found that gene networks with and without feedback motifs can exhibit differences at the level of how trait variance is partitioned and at the level of the number of apparent loci underlying trait variation. Although all gene network motifs were found to lead to statistical epistasis, positive feedback were found to do it much more that any other network motif. In addition, positive feedbacks and cyclic dynamics gave more non-monotone GPMs (i.e. maps where order is not preserved between allele content and corresponding genotypic value) and much lower heritabilities than those without.

In a recent study by Van Dooren (2018) found no prediction bias when applying the breeders equations to two traits in the eyespots of the butterfly. The currently accepted hypothesis for eyespot development is based on a gradient model for the concentration of a diffusible signal (Dilão and Sainhas, 2004). Cells in the center of the eyespot secrete a morphogen; the color and size of the eyespot is determined by thresholds to the concentration of the morphogen. The formation of the eyespot as we understand it can therefore be reduced to a one-dimensional process, without major cell rearrangements co-occurring with the patterning. This process is therefore unlikely to produce nonlinearities as large as the ones we find for cusp patterning in the tooth development model.

### How important are GPM nonlinearities in a population context?

It has been proposed that nonlinearities in the GPM are not relevant, at least in the short term. Quoting (Hansen 2008): “Sometimes biologists without statistical training will dismiss the additive model with the argument that genes exert their effects through highly nonlinear physiological interactions. It is almost guaranteed, however, that the additive model will be a good local approximation to the GPM for segregating genotypes. This is due to the statistical definition of gene effects as averages over the genotypic combinations in which they occur. Even if the effect of a particular gene substitution may be different in different specific genetic backgrounds, the averaging over all backgrounds minimizes the variation.”. Although it is true that such minimization occurs, there is no reason to assume it would necessarily be strong enough to make the nonlinearity of the GPM irrelevant for short term predictions. This depends on how strong such nonlinearities are and, to a lesser extent, on the population size and recombination rates (see section *The effect of populaton parameters on the bias* below). In fact, our results explicitly show that this is not the case, even with large populations in linkage equilibrium (see Fig. S12). The GPM is often nonlinear enough to lead to significant deviations between observed changes and predicted changes in the short term.

To understand, from a quantitative genetics perspective, why breeder’s equations may still fail despite the averaging over genetic backgrounds, we must distinguish the functional, individual-level description of the GPM and the statistical, population-level one used in quantitative genetics. The marginal parameter-phenotype maps shown in our work (Fig. 3) are made at the individual level: interactions at the molecular, cellular and tissue level that result in individual variation. It is independent of the allelic variants present in the population, and purely determined by the developmental and genetic system behind the development model. The parameter-phenotype map at the population level is evidently related to the individual-level one (Alvarez Castro and Carlborg 2007), but it is weighted by the allelic frequencies existing in the population. At the population level, the map describes how a change in parametric values result in an average change in the mean phenotypic value in the population. Since in our model the mapping between loci and developmental parameter values is additive, we can assume that the parameter values are distributed normally in the population. The population-level parameter-phenotype map can then be thought of as the individual-level map passed through a Gaussian filter, as shown in Fig. S12. Such smoothing reduces the ruggedness compared to the individual-level map, and does so depending on the spread of the parameter values in the population (a proxy for allelic distributions). This population-level smoothing can be enough for the map to be adequately described locally by a plane, leading to accurate description by multivariate breeder’s equation. This will be the case if the individual-level map is already linear (as in Fig. S12A) and even sometimes when it is nonlinear (as in Fig. S12C). As we show in this work however, the smoothing may not be enough for a relatively high percentage of generations when large nonlinearities at the individual level result in large nonlinearities at the population level (as in Fig. S12E). In terms of average genetic effects, this means that certain nonlinearities at the individual-level will result in gene substitutions with effects that are highly dependent on the genetic background. For a given gene substitution we can think of an associated distribution of genetic effects across the different backgrounds. The shape of this distribution will depend on both the substitution itself and the genetic backgrounds present in the population. When reducing this distribution of effects to its mean value – as when calculating the average genetic effect – we end up with a statistic that poorly represents the distribution of effects. This ultimately leads to predictions based on additive effects being inaccurate, even for single-generation predictions.

### The effect of populaton parameters on the bias

Prediction bias was found across all population genetic parameter sets studied (see Materials and Methods and Fig. 4). In general, our exploration shows a saturating relationship between the evolutionary parameters and the prediction bias. This is, there is some dependency of the bias with the parameters but the amount of bias reaches a plateau, determined by the characteristic of the GPM which are not dependent of population-level parameters. The characteristics of the GPM are determined by development only, the population genetics parameters affect bias indirectly by affecting how fast the population moves in the developmental parameter space and, thus, the likelihood of finding a region of such space with a complex GPM.

The effect of selection strength was studied by deviating the factor in the exponent of the selection function. In each generation, the fitness of each individual in the population was calculated as a function of the distance of its trait values and the optimal trait values as

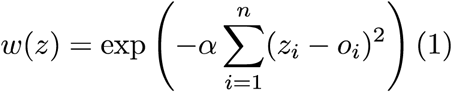

Where *z* is the vector of trait values for the individual, z_*i*_ is the i^*th*^ element of that vector, o_*i*_ is the i^*th*^ element of the vector of optimum trait values and is the fitness function. Therefore, the closer an individual is to the optimum morphology, the higher its fitness. The steepness of the selection gradient is determined by the parameter *α* (see Materials and Methods). Stronger selection makes the population move faster in the developmental parameter space. This allows for it to explore larger areas of the developmental parameter space and therefore increasing the chances of passing through highly nonlinear regions of it.

Mutation rate is the per-allele probability of being mutated. Larger mutation rates lead to larger percentages of significant bias. This is explained by the fact that more mutations lead to larger exploration of the GP map and bigger probability of reaching problematic regions. The core and low mutation rates do not show large differences.

Mutation strength determines how large is the phenotypic effect of a mutation in respect to a reference value. Tinkering this parameter leads to a a similar plateau-like response in the amount of significant bias as seen when tinkering the mutation rate.

Larger populations show larger percentage of significant bias. This can be explained by two facts. First, larger populations explore larger proportions of the developmental parameter space, and therefore they have larger chances of encountering nonlinear regions. On the other hand, smaller populations experiment larger drift effects and it therefore becomes harder to confidently say that the prediction is different to the observed change in our repetition experiments. In this latter case, we detect bias less often because there is less statistical power to detect it.

Finally, there is not a clear dependency between the amount of bias and the number of loci additively determining each parameter. This is due to two counteracting effects. On the one hand, with a fixed per-allele mutational rate, more loci imply more mutations per genome. On the other hand, since more loci imply that each of them contributes less to the parameter value, fixed mutational strength means each mutation is smaller. This means that when increasing the number of loci determining each parameter, there are more mutations of smaller effect, and eventually to no big changes in the amount of bias detected.

## III Supplementary tables

**Table S1.** Parameter values for the simulations. The middle columns shows the core population parameter set. Simulations with all optima were ran by changing one of the population parameters above at a time. This leads to a total of 11 population parameter combinations ran, and a total of 220 simulations.

|  |  |  |  |
| --- | --- | --- | --- |
| Population size (N_pob) | 100 | 600 | 1000 |
| Number of haploid loci (N_loci) | 42 | 105 | 210 |
| Mutation standard deviation (tau) | 0.02 | 0.1 | 0.5 |
| Allelic mutation rate (muta) | 0.01 | 0.001 | 0.0001 |
| Selection strength (alpha) | 0.125 | 0.05 | 0.01 |

## IV Supplementary Figures

**Fig. S1:**
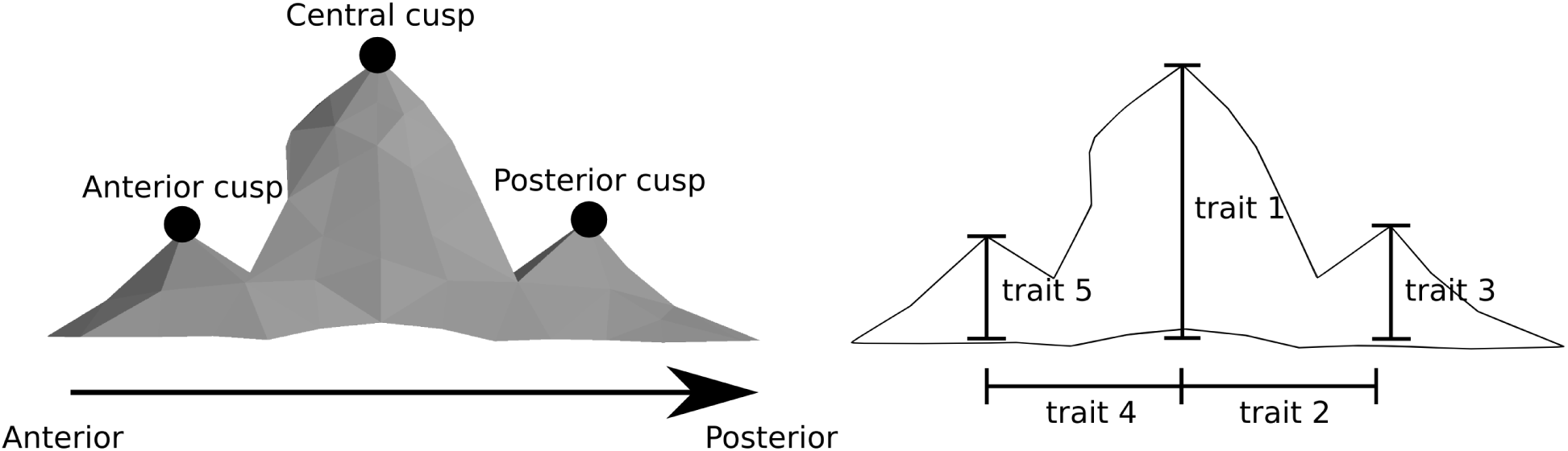
Example 3D morphology arising from the developmental model. In the left we mark (black balls) the landmarks used to define the traits (right) on which natural selection is applied. The height, y-coordinate of the landmarks, is calculated as the difference from the mean y position of the cells in the tooth margin (as in ^16^).

**Fig. S2:**
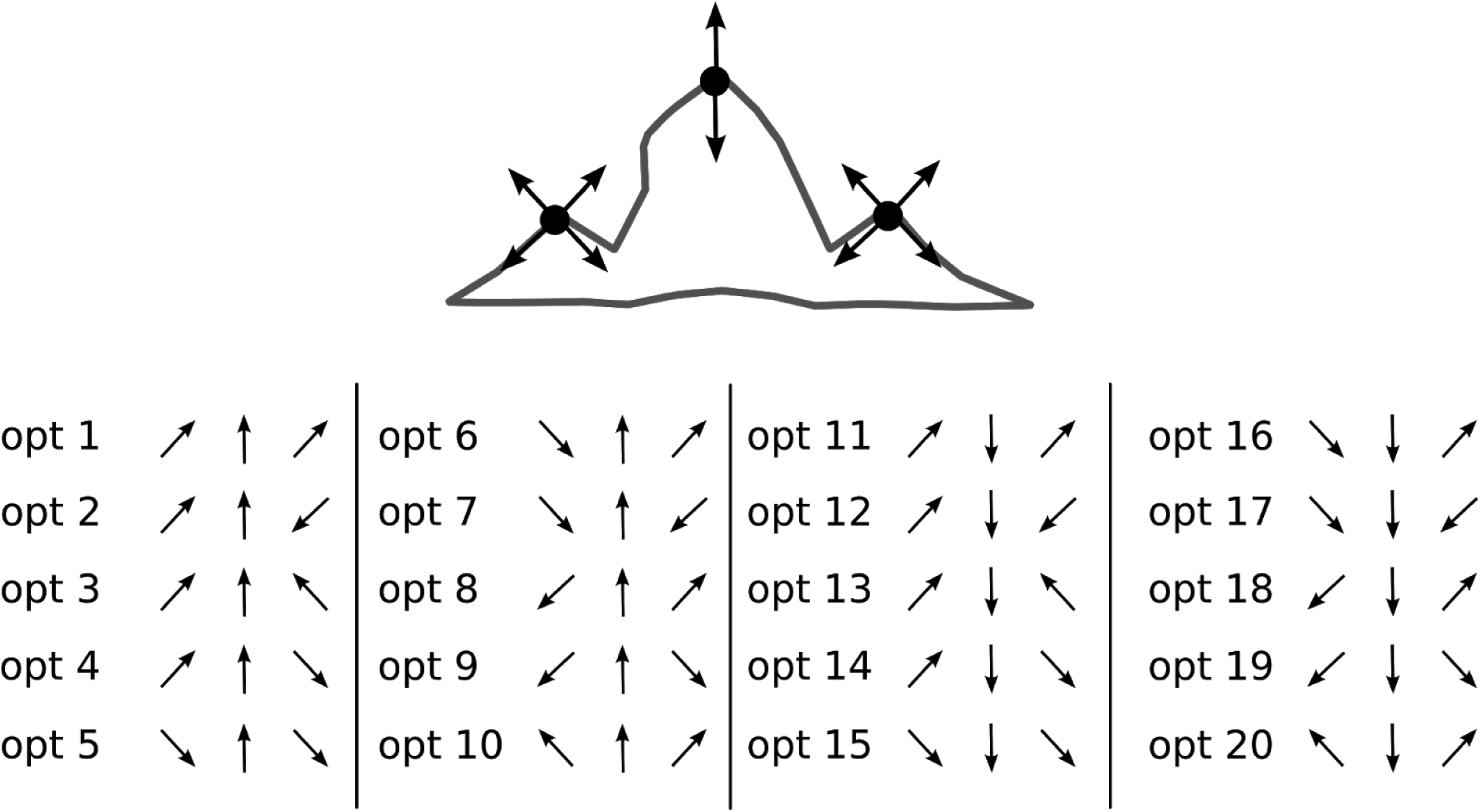
List of optima used in the simulations. Optima are shown as deviations from the initial morphology, with arrows symbolizing the direction of selection for each of the three cusps. Each of the 5 traits defined in Fig. S1 can be selected to increase or decrease, leading to a total of 2^5^ = 32 trait combinations. We have removed the ones are mirror images of others. Because we wanted selection to act with the same strength on every trait, the optimum value for each trait was set at an equal distance from its mean value in the starting population. The optimum morphology was then determined by adding or subtracting 3 units to the mean value for the trait in the population at generation 1, depending if selection was upwards or downwards, respectively.

**Fig. S3:**
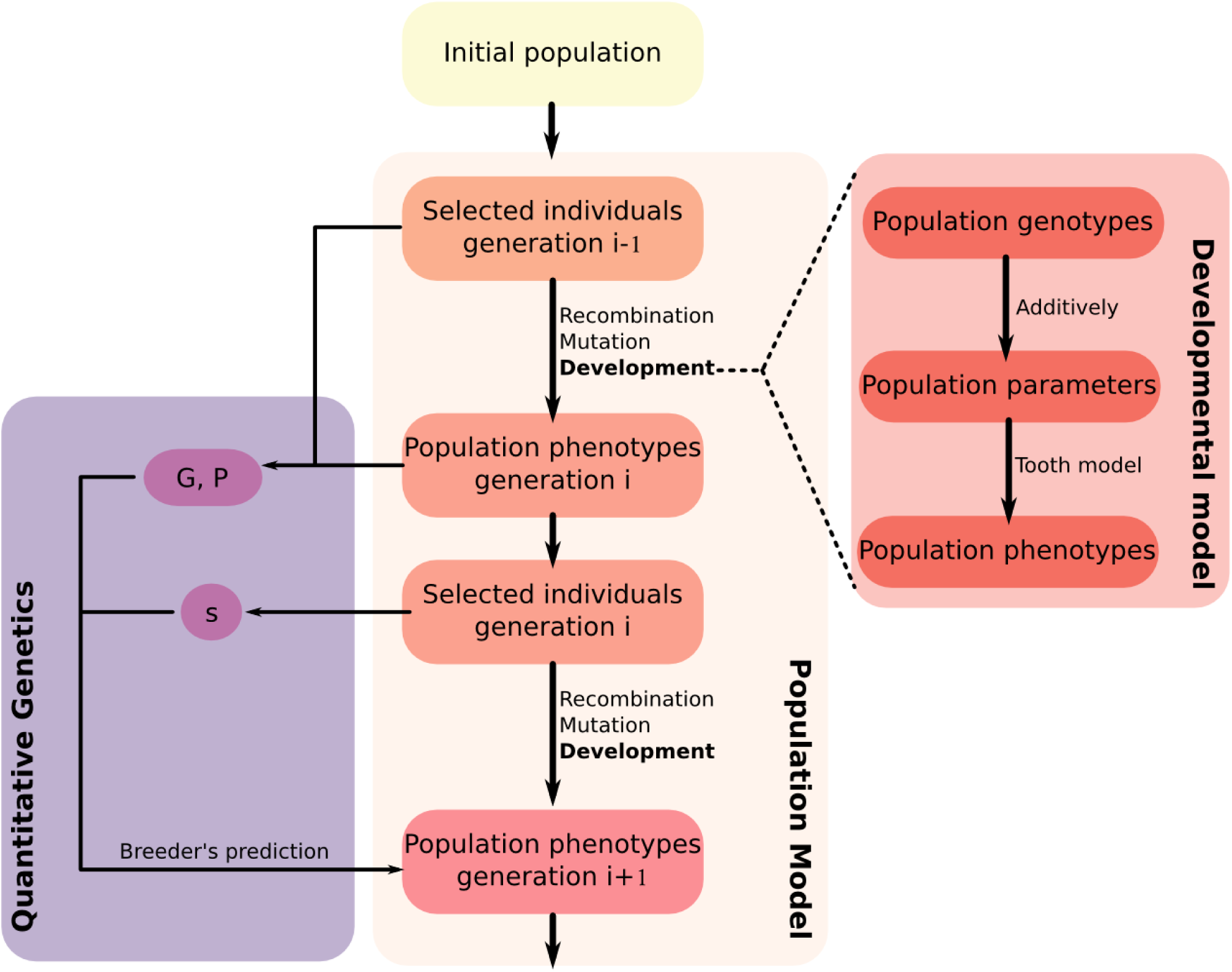
Evolutionary algorithm. The scheme shows the steps of the population model algorithm used in our simulations. Each step is explained in the text.

**Fig. S4:**
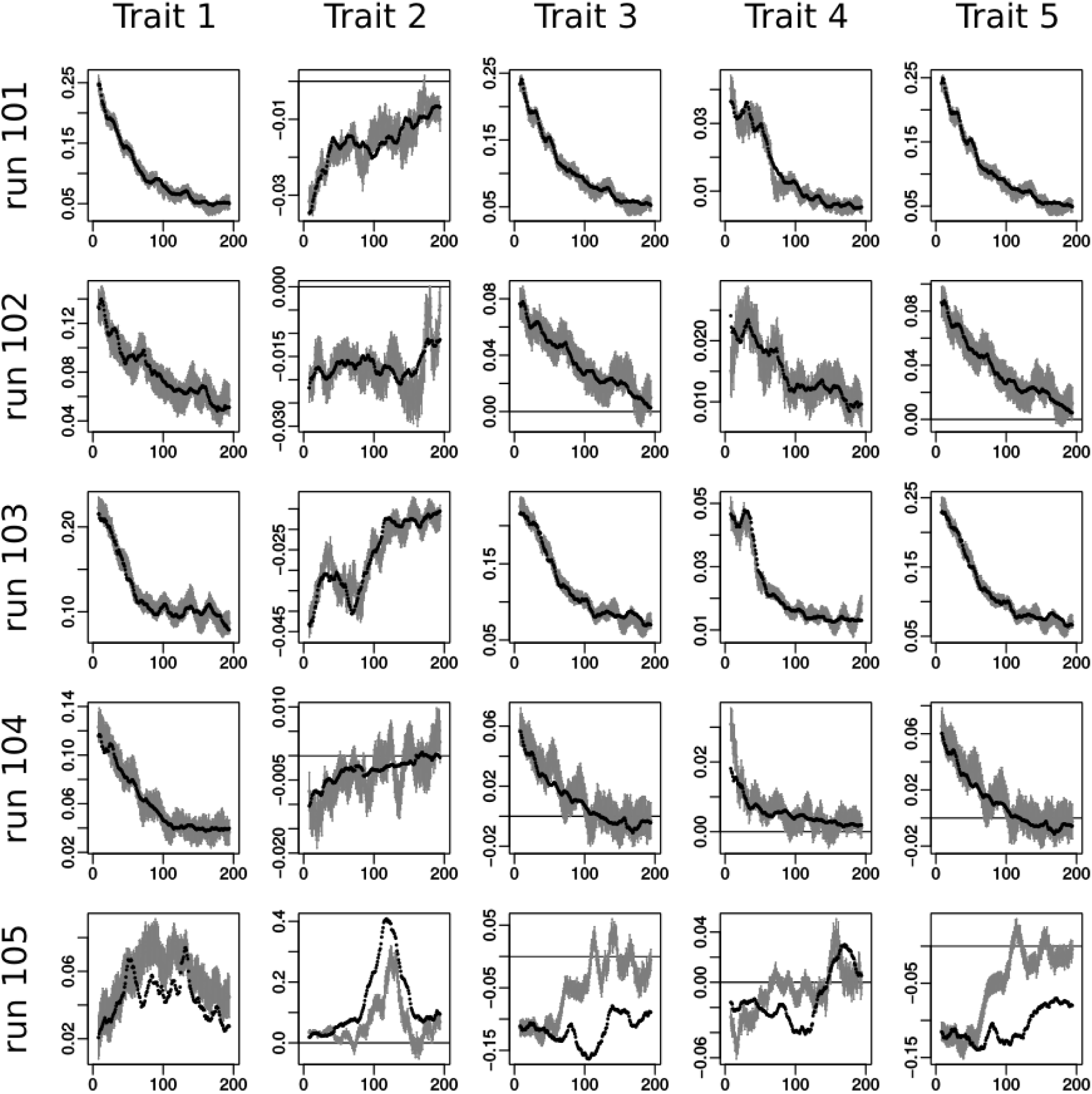
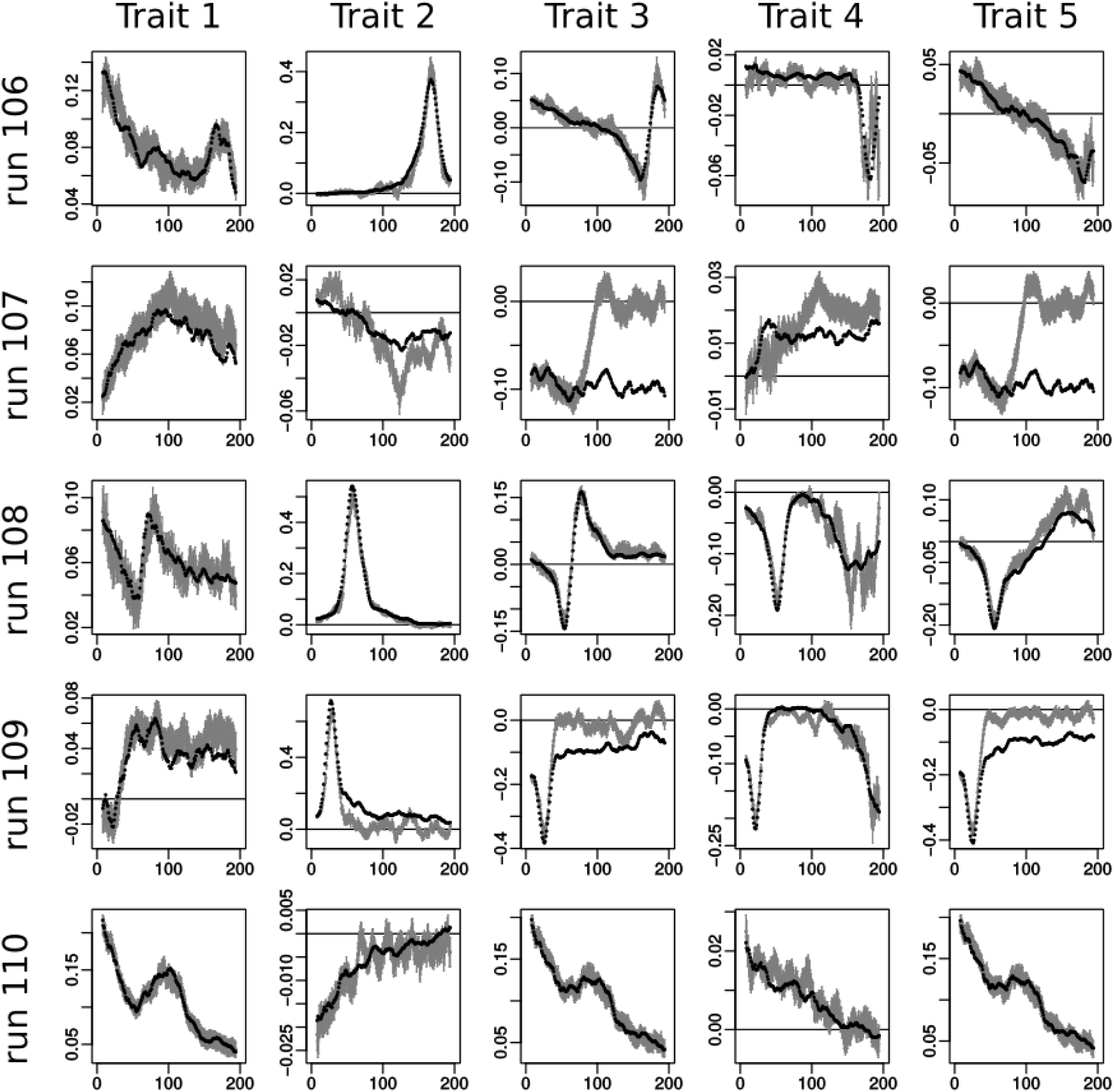
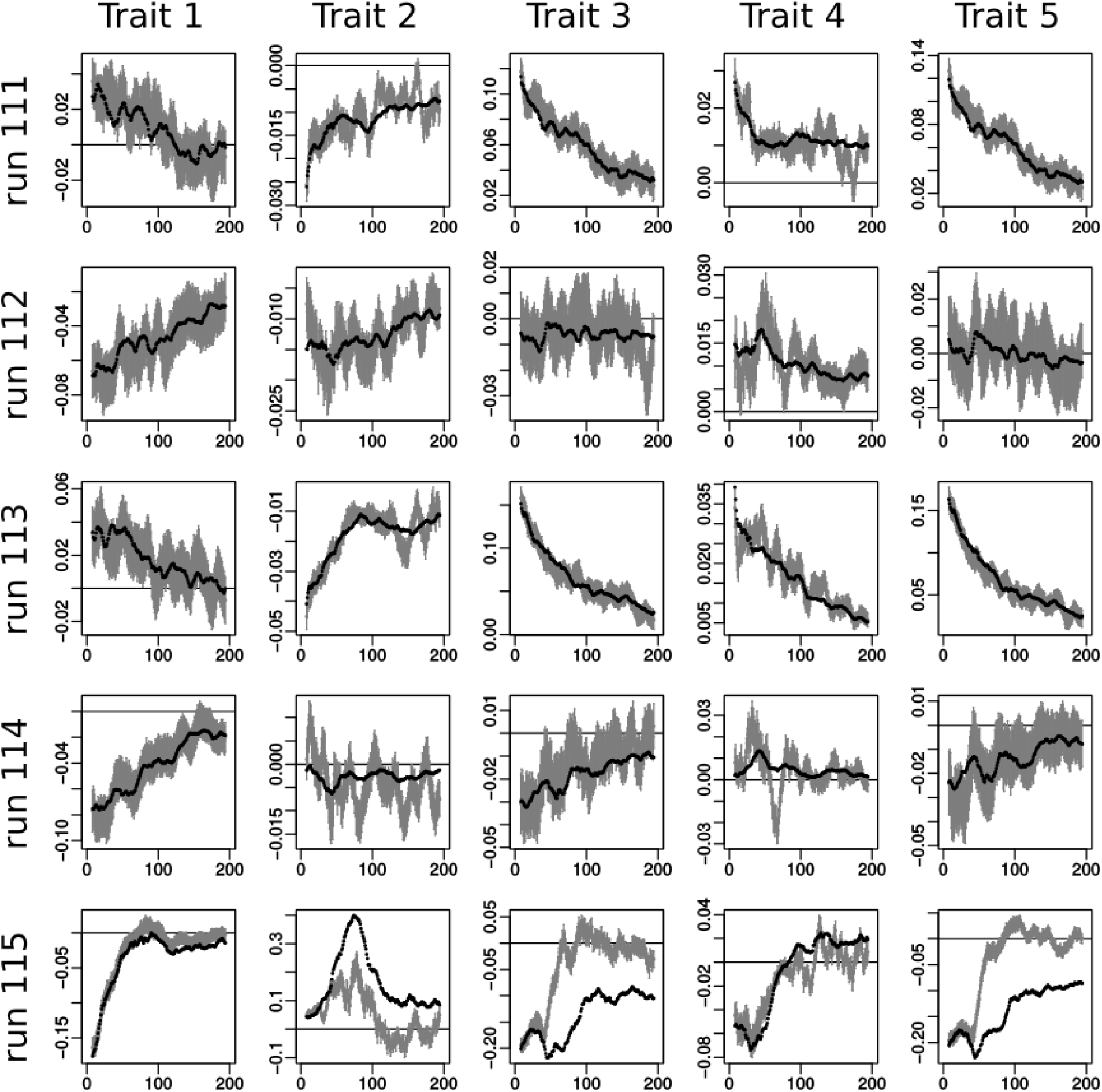
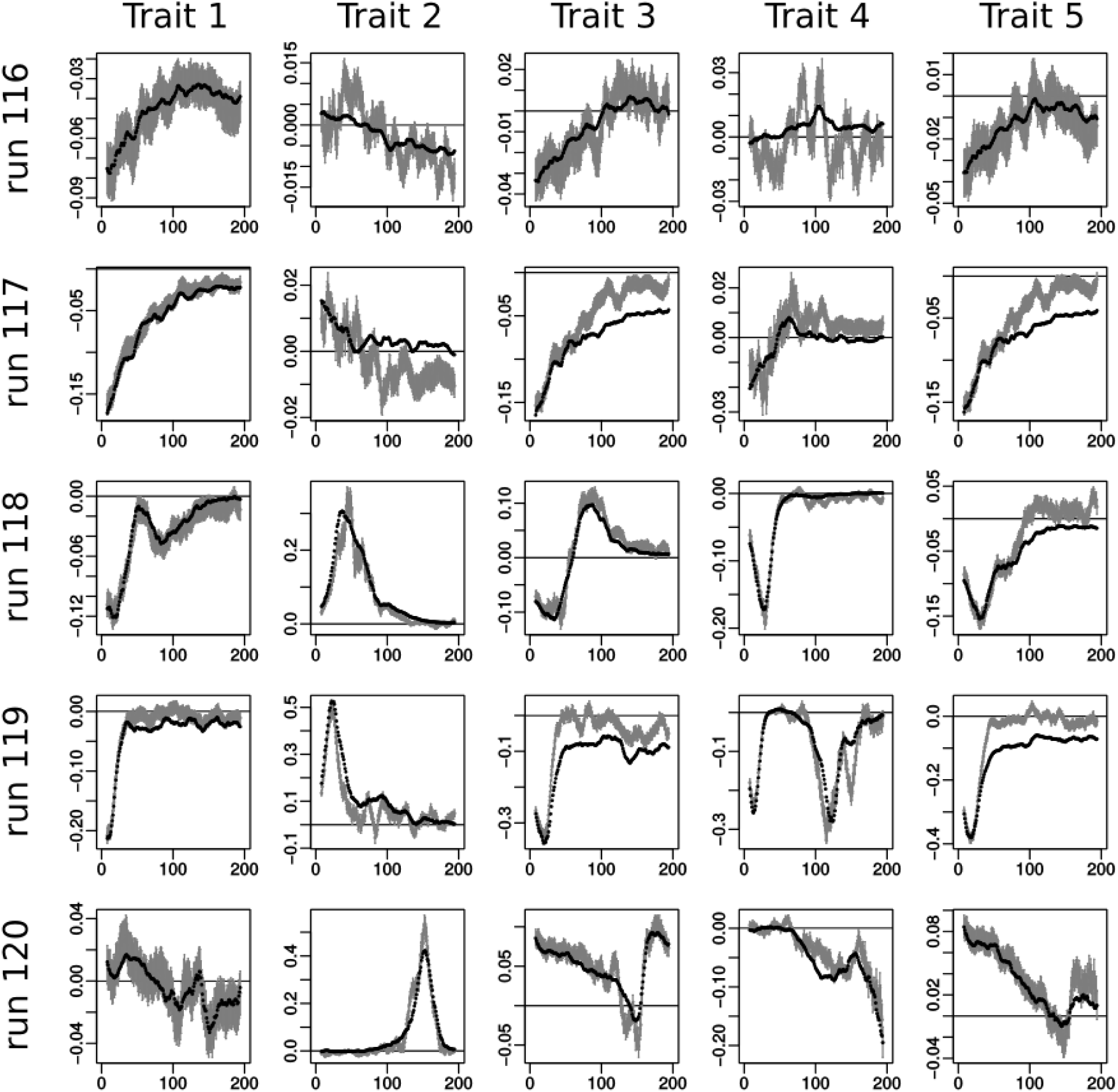
Observed and predicted change can differ in a systematic way over generations. As figure 1 but for all simulations with the 20 different optima in the core parameter set (see Table S1).

**Fig. S5:**
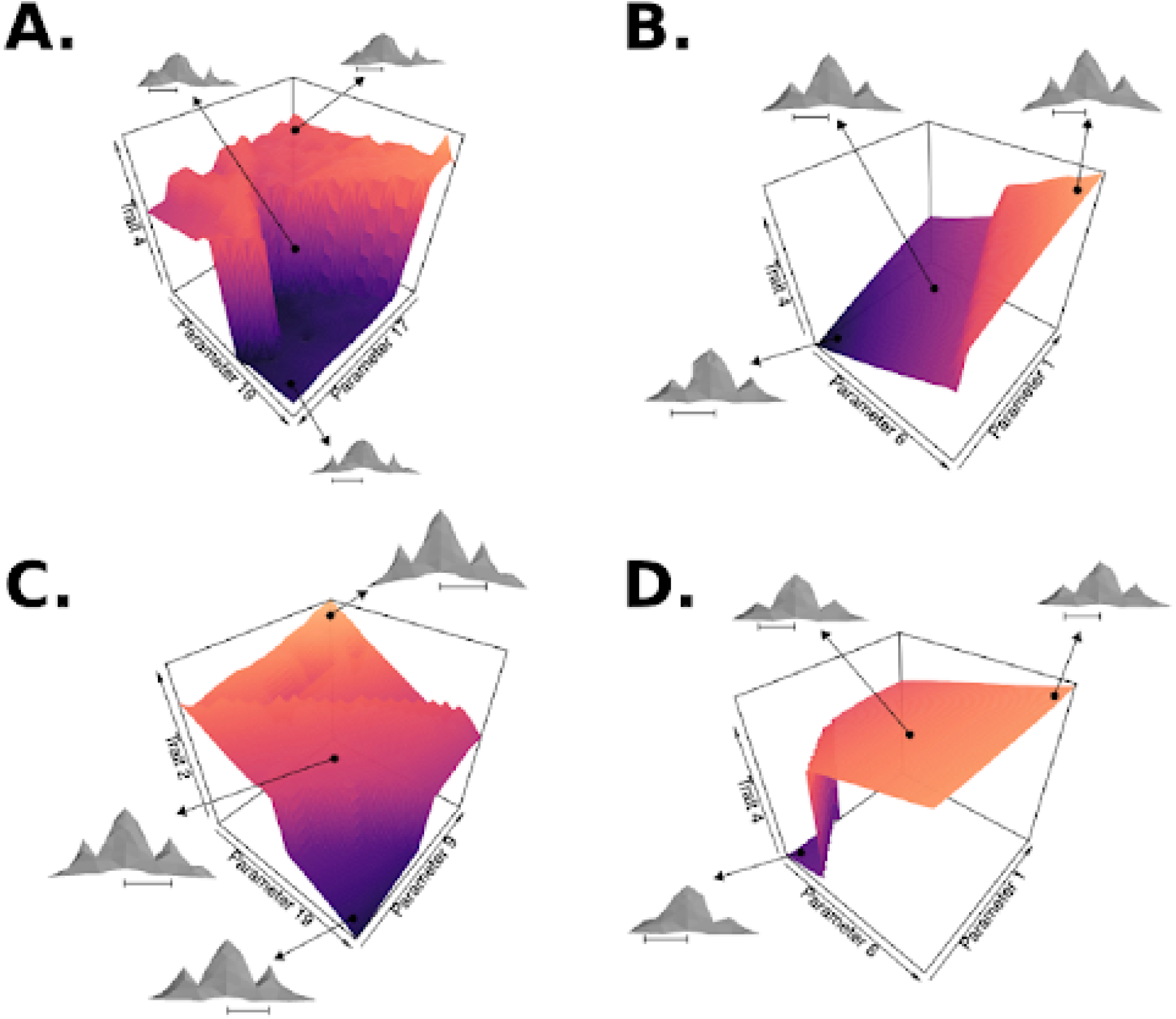
Prediction bias arises when the population is in a nonlinear region of the parameter-phenotype map. As Fig. 3 but for other examples.

**Fig. S6.**
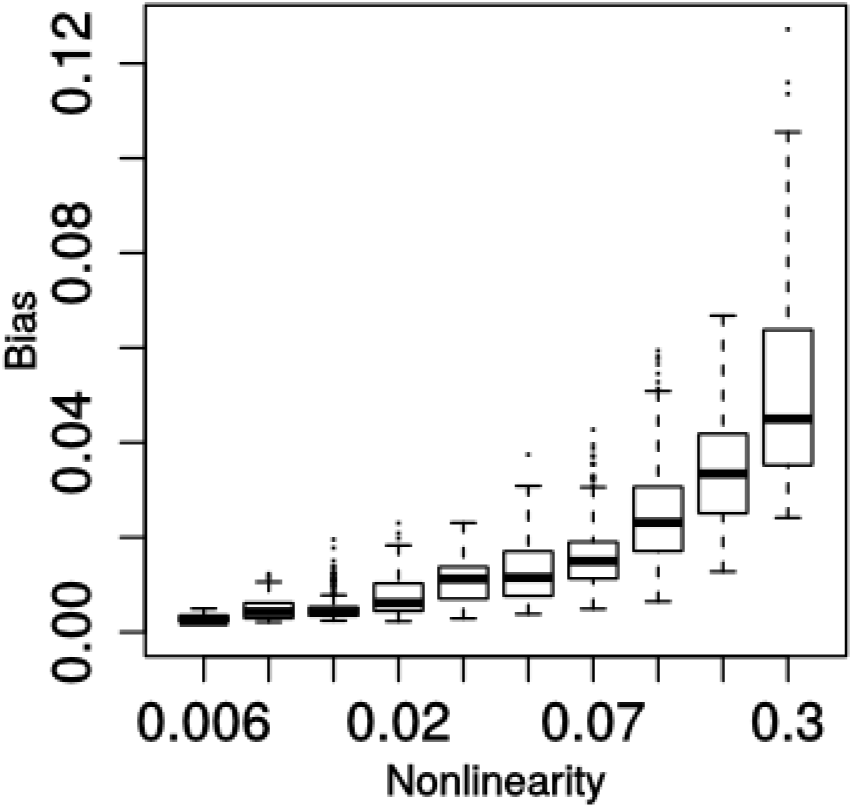
Prediction bias correlates with the local non-linearity of the genotype-phenotype map. Absolute bias versus local nonlinearity of the parameter-phenotype map for all simulated generations showing a significant bias (see Fig. 2). Both the frequencyspread and magnitude of the bias increase with the nonlinearity of the parameter-phenotype map. Each point represents the bias and nonlinearity in a trait, in a generation of a simulation in the core set. Points were grouped into bins and a boxplot was made for each bin. Local nonlinearity was measured based on how well a linear map describes the relationship between parameters and traits. A best linear fit (*B* from the vector of mean developmental parameters in the population (*u*) to the vector of observed traits (*z*) is found by least squares, using the data from all individuals in the population in that generation: **B** = **u**^*T*^(**uu**^*T*^)^-1^ Where T is the transpose operator. Both phenotype and trait values are expressed as deviations from the mean. The root-mean-square error of the fit is taken as a measure of nonlinearity for each trait, i.e. 0 means the map is perfectly described by the linear fit.

**Fig. S7:**
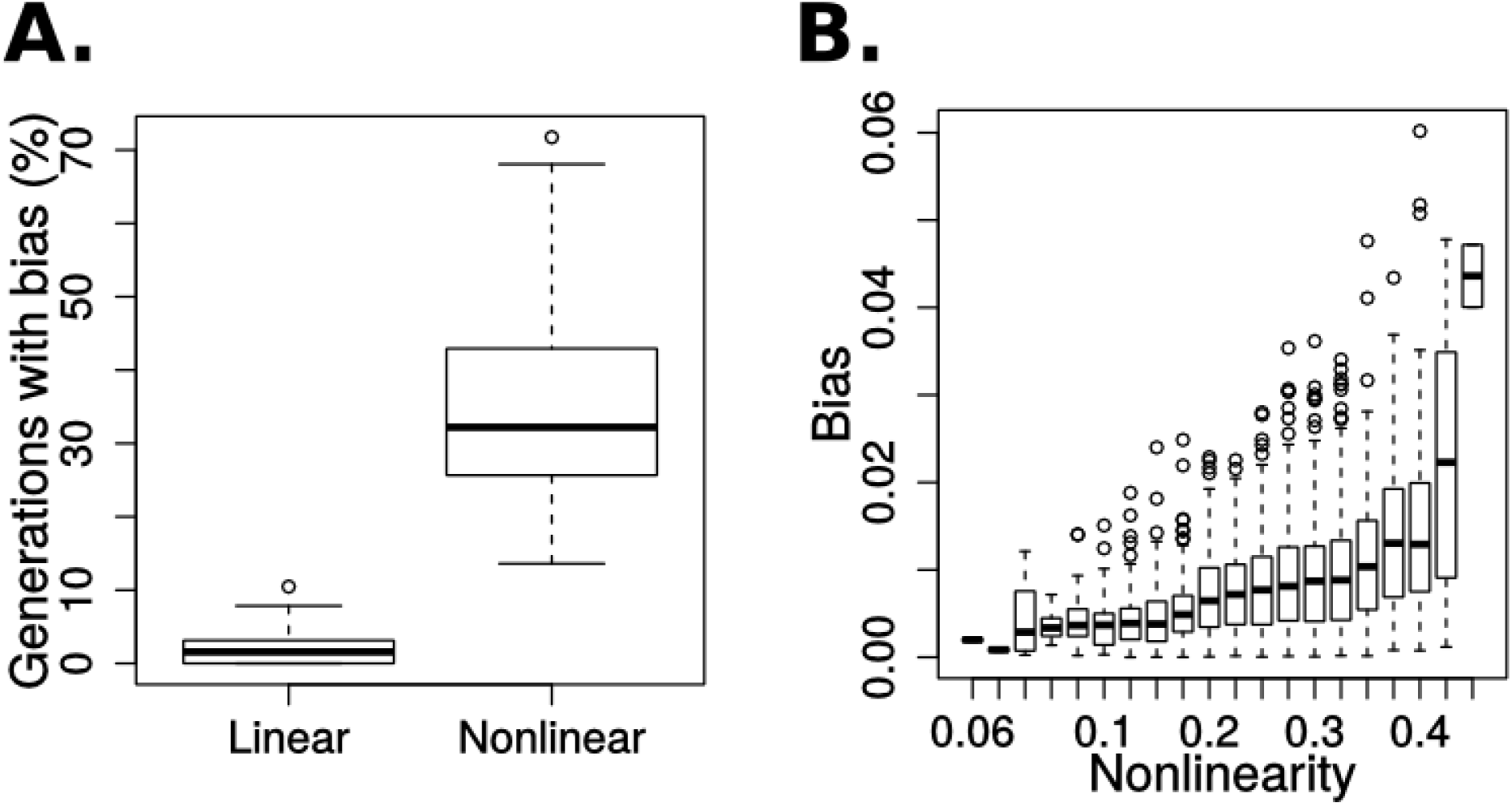
Bias is common for any non-linear GPM. The plots shows the percentage of generations with significant bias and a binned scatter-plot of local non-linearity of the parameter-phenotype map against prediction bias for two analytical GPMs. **A** shows the distribution of percentage of generations with significant bias (99%) for all 30 simulations ousing the linear and the nonlinear GPMs. When the GPM is linear there is negligible amount of detected bias (median of 1.57% of generations). When the GPM is nonlinear, prediction bias is very common. **B** shows, for the non-linear map, the absolute prediction bias against the local nonlinearity of the parameter-phenotype map as shown in Fig. S6. Both the spread and mean amount of bias increases the nonlinearity of the parameter-phenotype map.

**Fig. S8:**
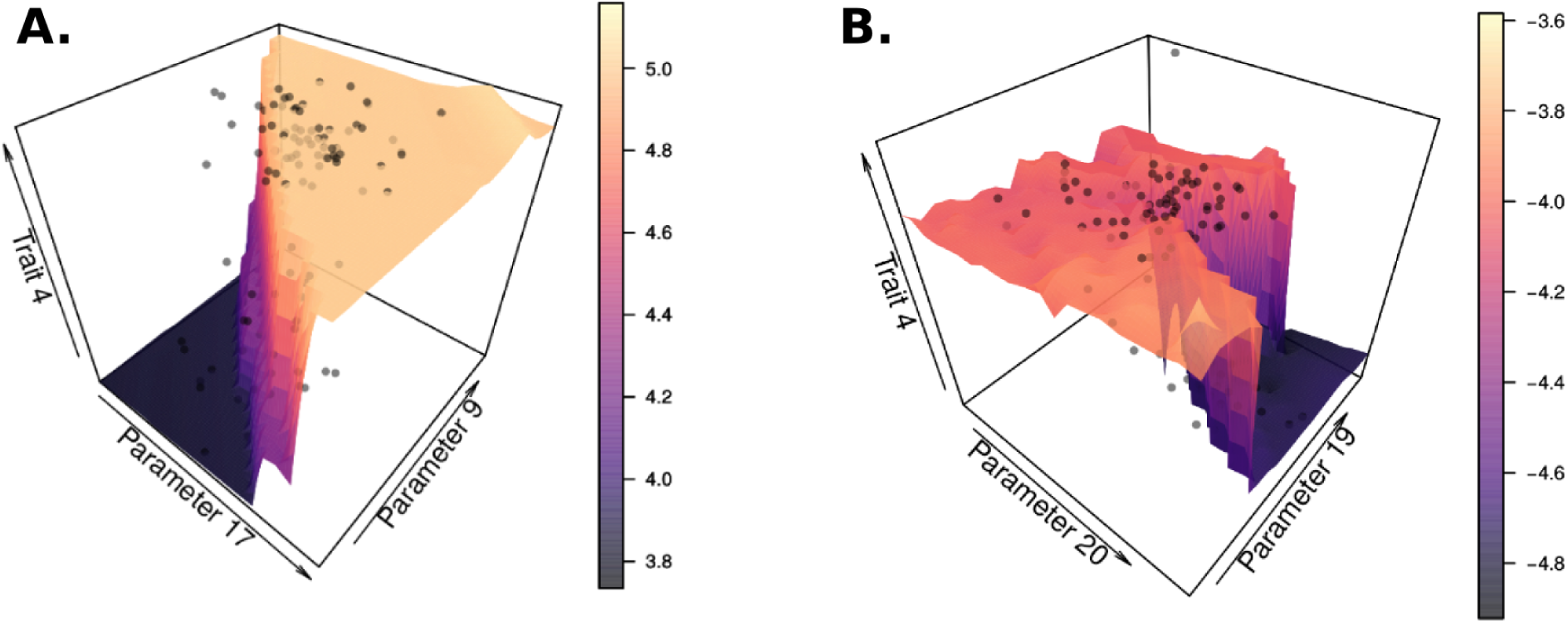
The relationship between parents and offspring trait values is not always linear, the population is distributed over a region of the parameter space that leads to different trait values in a non-gradual way: Marginal parameter-phenotype maps including 100 individuals in the population as black points. Individuals do not always fall on the surface of the plot because of variation in parameters that have not been plotted (that is why we call these maps marginal).

**Fig. S9:**
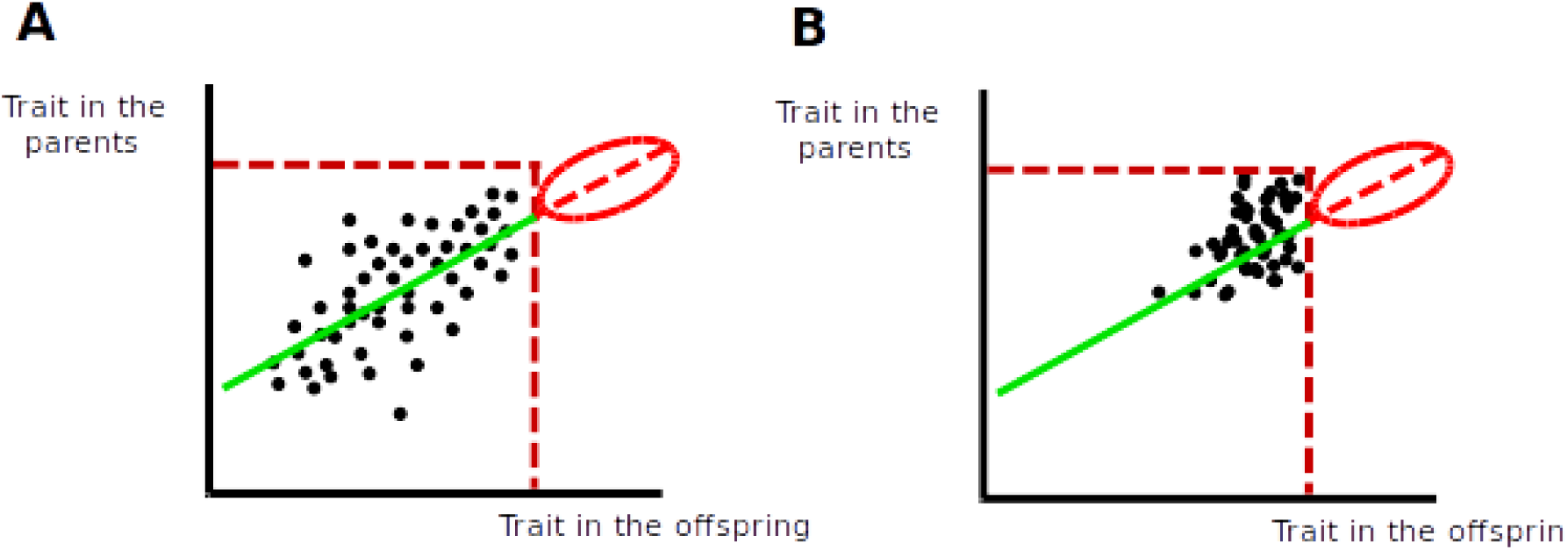
Bias can arise because variation does not exist beyond a trait value but the linear transformation in the breeder’s equations infers that such variation should exist. Idealized univariate example of how a limitation in a trait value leads to bias in the breeder’s predictions. An actual example can be seen in the dashed line of Figure 3G. This example allows to more clearly explain one way in which bias arises. **A** and **B** show the parent-offspring regression of a trait (although the same could occur between traits). The dashed vertical and horizontal red lines show the value over which a trait can not pass due to developmental dynamics. Simply the population is in a region of the developmental parameter space where this trait is always below this value in all possible phenotypes. Regions of that space that are far away from where the population is now may be able to produce larger values in this trait. The green line shows the regression between parents and offspring. From this regression one would infer that trait values over the dashed vertical red line should be possible while they are not, red ellipsoid. **B** shows the same that **A** but for a population that has encountered this bias long ago and in which natural selection favours large values in the trait. Natural selection pushes the population towards the dashed vertical red line but due to the always present mutation and recombination there are always individuals with lower values of the trait. Because of that not all individuals are on the limit of the possible values (dashed vertical red line) and a non-zero regression exist. Because of this regression breeder’s prediction is still that the trait should increase beyond the non-possible trait value, thus, the bias.

**Fig. S10:**
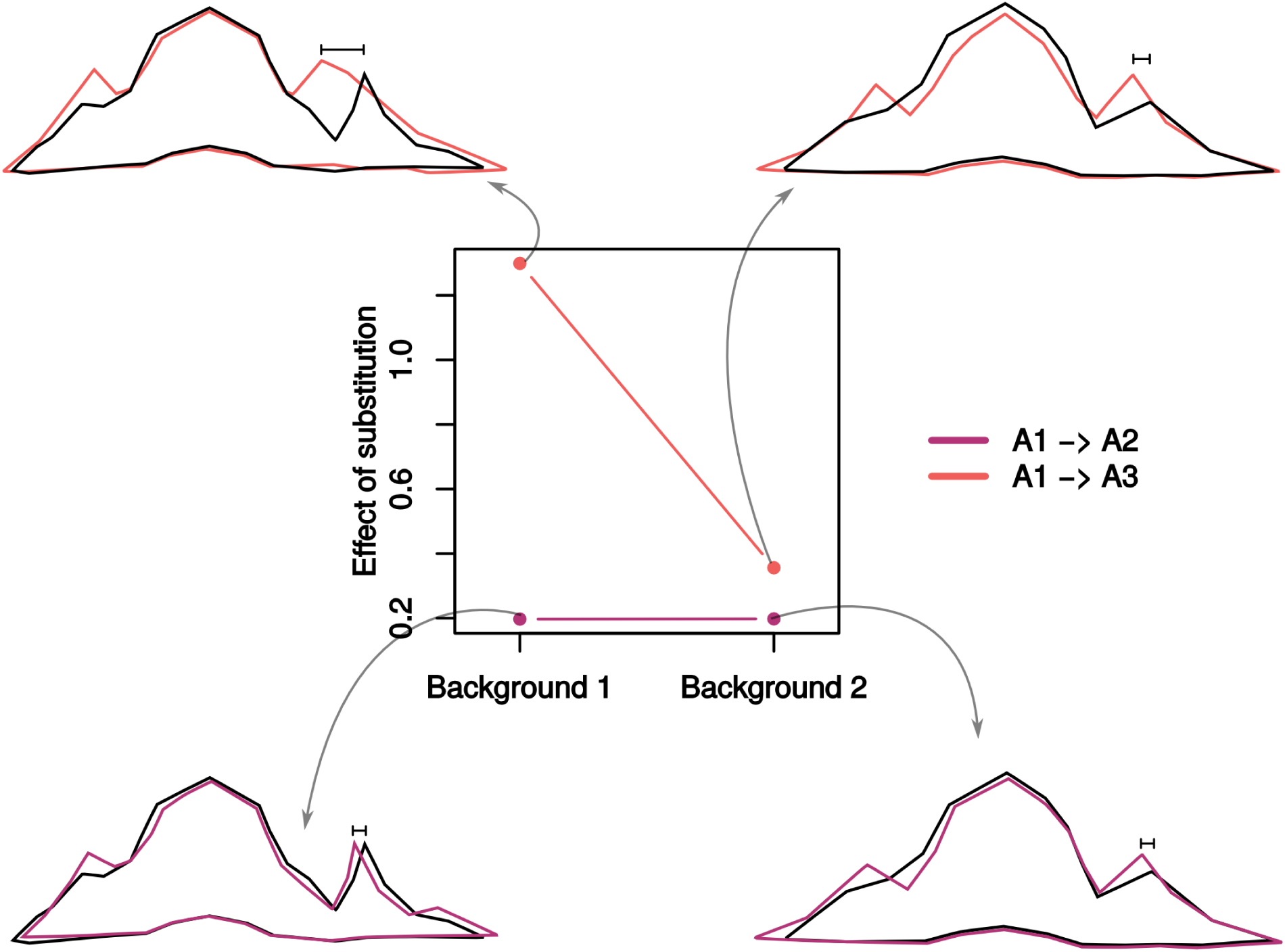
The tooth model is not compatible with the assumptions of the multilinear model. The multilinear model developed by Hansen and Wagner^21^ assumes that changing the genetic background for a locus causes a linear transformation of the effects of all substitutions in that locus. The figure shows an example where this does not hold using the tooth model: two allelic substitutions are scaled by different factors when changing the genetic backgrounds. In the example we measure trait 2, the x-position of the posterior cusp. We quantify the difference in the value of this trait when making two allele substitution, with respect to the phenotype of the reference allele (A1). Included in the plot are outlines of the teeth morphologies. Outlines in black correspond to the reference morphologies. Colored outlines correspond to the morphologies obtained after the allelic substitutions, in the two backgrounds. A black segment indicates the difference in the trait value between the reference morphology and the one produced after the allelic substitution, which is also the y-axis in the plot. The allelic substitution A1->A2 has almost the same phenotypic effect in both backgrounds. The substitution of A1->A3 however, produces a much larger effect in Background 1 than in Background 2, violating the assumptions of the multilinear model. We denote 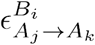 as the genetic effect of substituting allele j (A_*j*_)with allele k (A_*k*_) on genetic background i (B_*i*_). For a reference allele A_1_ and for two genetic backgrounds B_1_ and B_2_, the multilinear model assumes that there exists a linear transformation T such that for every allele A_*k*_, 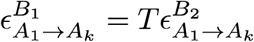. Equivalently, it holds that for every A_*k*_,

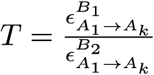 Assuming

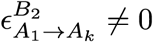 In the example, this does not hold,

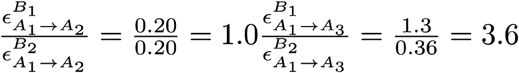

**Fig. S11.**
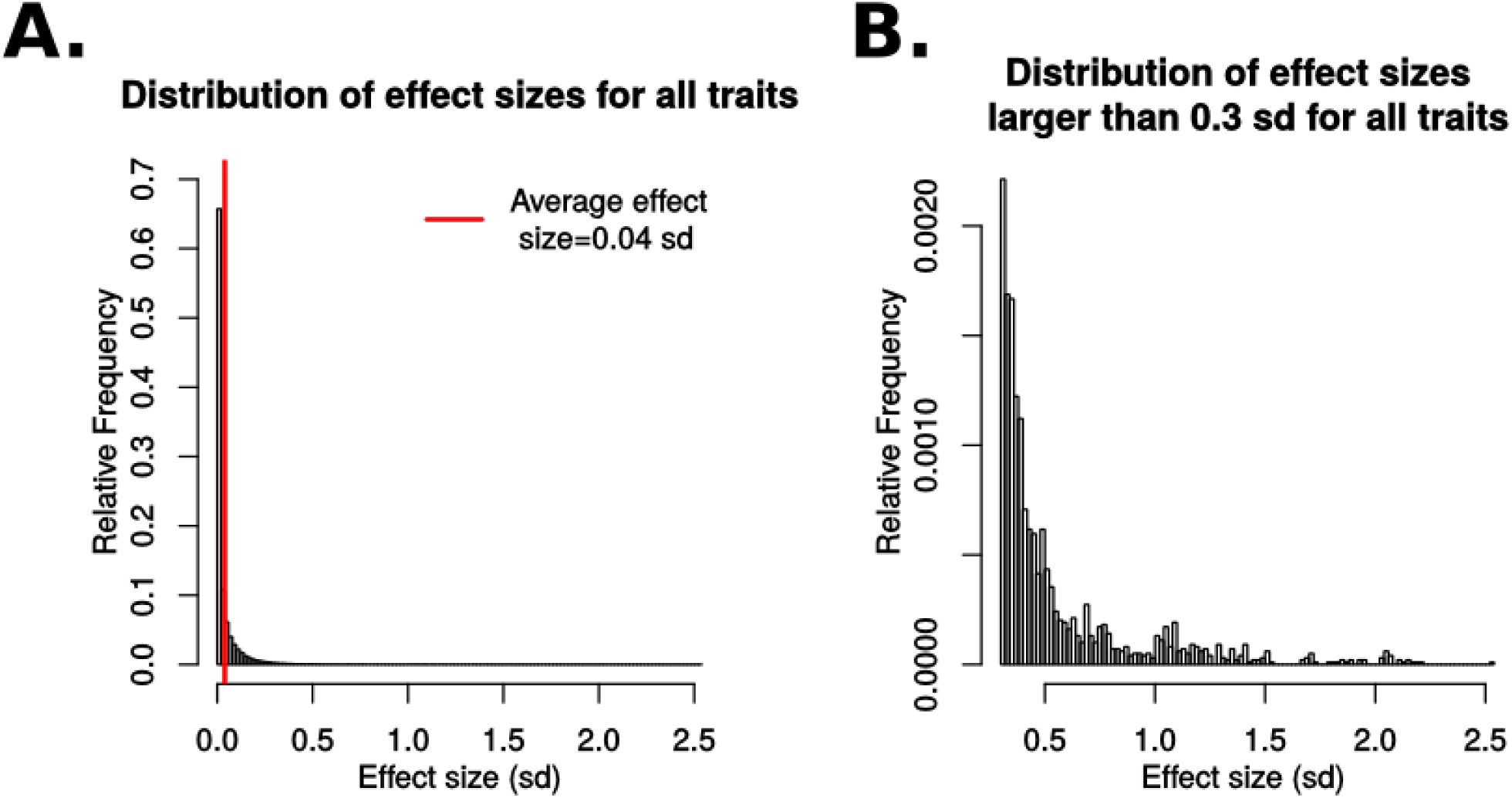
The effect size of mutations falls within a realistic range. Distribution of the effect size of mutation on traits relative to trait’s standard deviation (for all 20 simulations in the core parameter set and sampled every 20 generations, starting from generation 10 to generation 190). First, the average value for each loci was calculated by averaging over the values present in the population. This results in a list of mean loci values, for all loci in the population. This list corresponds to the “reference individual”, and its corresponding phenotype is calculated by running the tooth development model. From the genotype of the reference individual, each loci is mutated, one at a time, with a mutation strength equal to the value in the core parameter set. This creates a set of single-loci mutants. The phenotype of each mutant is generated using the tooth development model. Finally, the deviation from each mutant to the reference individual is measured for each trait, and divided by the standard deviation for that trait in the population. Panel A shows the distribution of these effects for all traits, including all mutants generated for all simulation and generations sampled. The mean effect is shown with a red line, and equals 0.04 standard deviations. Because most effects are very small, larger effects cannot be seen from the plot. Panel B shows only the effects that are larger than 0.3 sd, which can be considered large effects. The distribution found is compatible with others previously reported QTL distributions, so the mutation strength in the core parameter set is not unrealistic. Note that our estimate for mean effect is close to 0 because we can detect even the smallest effects, which is impossible to do with real-life data. The data fits the expectation of an exponential-like distribution of effects sizes.

**Fig. S12:**
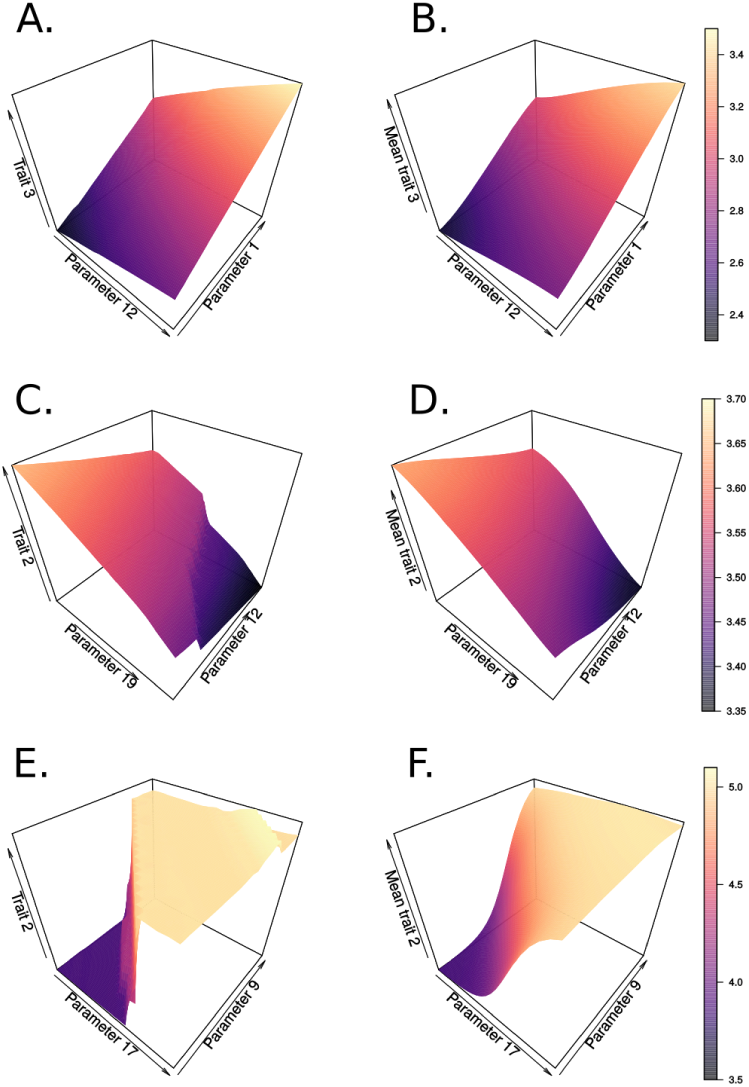
Individual-and population-level parameter-phenotype maps. Individual-level parameter-phenotype maps (A, C and E) were constructed to see the phenotypic effects of changing two parameters while keeping all the others constant. Here, all-but-two parameters were fixed to their mean value in the population at the generation plotted. The remaining parameter was varied around their mean values and within the limits of values existing in the population. The trait values of the resulting morphologies were measured and plotted as the z axis. Population-level maps (B, D and F) were constructed by running a Gaussian smoother through the individual-level map. These maps therefore indicate the average trait value for an average parameter value. We assume a Gaussian distribution of parameter values in the population. The plot shows that if the individual-level map is linear (A), the population-level one will be as well (B). If the individual-level map is nonlinear however (C and E), the population-level map may or may not show significant nonlinearities (D and F), depending on the magnitude of the individual-level nonlinearities and the spread of the parameters in the population.

